# Structure and function of yeast Lso2 and human CCDC124 bound to hibernating ribosomes

**DOI:** 10.1101/2020.02.12.944066

**Authors:** Jennifer N. Wells, Robert Buschauer, Timur Mackens-Kiani, Katharina Best, Hanna Kratzat, Otto Berninghausen, Thomas Becker, Wendy Gilbert, Jingdong Cheng, Roland Beckmann

## Abstract

Cells adjust to nutrient deprivation by reversible translational shut down. This is accompanied by maintaining inactive ribosomes in a hibernation state, where they are bound by proteins with inhibitory and protective functions. In eukaryotes, such a function was attributed to Stm1 (SERBP1 in mammals), and recently Lso2 (CCDC124 in mammals) was found to be involved in translational recovery after starvation from stationary phase. Here, we present cryo-electron microscopy (cryo-EM) structures of translationally inactive yeast and human ribosomes. We found Lso2/CCDC124 accumulating on idle ribosomes in the non-unrotated state, in contrast to Stm1/SERBP1-bound ribosomes, which display a rotated state. Lso2/CCDC124 bridges the decoding sites of the small with the GTPase-activating center of the large subunit. This position allows accommodation of the Dom34-dependent ribosome recycling system, which splits Lso2-containing but not Stm1-containing ribosomes. We propose a model in which Lso2 facilitates rapid translation reactivation by stabilizing the recycling-competent state of inactive ribosomes.

## Introduction

Ribosomes are the universally conserved biological machines which translate genetic information from mRNA templates into polypeptides with the corresponding amino acid sequence. Maintaining ribosome functionality is vital for cell survival under all circumstances. Hence, regulatory mechanisms have evolved which facilitate transition of the translation machinery between active and dormant states. This allows cells to dynamically adapt to environmental changes in nutrient availability.

Nutrient starvation induced cellular stress and reversible translational repression is particularly well studied in prokaryotes. In bacteria, several small ribosomal binding factors (RBFs) have been identified which inhibit translation and facilitate reversible formation of inactive 100S ribosome dimers (Basu and Yap, 2017; Ortiz et al., 2010). These 100S dimers allow ribosomes to enter a hibernation state during stationary growth or stress phases (Beckert et al., 2017; Beckert et al., 2018; Matzov et al., 2017). Within hibernating bacterial ribosomes, RBFs (RMF, HPF, RaiA) bind the decoding center, occupy the mRNA binding channel, and block A and P tRNA sites (Ueta et al., 2008; Ueta et al., 2005; Yoshida et al., 2002). As a result, bacterial hibernation factors protect the crucial active sites of the ribosome and inhibit binding of both mRNA and tRNA, blocking translation altogether (Prossliner et al., 2018).

Several types of inactive or hibernating ribosomes have also been observed in eukaryotes (Brown et al., 2018; Krokowski et al., 2011). In general, idle 80S ribosomes accumulate after exposure to various stresses like amino acid shortage (Krokowski et al., 2011; Tzamarias et al., 1989), osmotic stress (Uesono and Toh, 2002) or glucose starvation (Ashe et al., 2000; van den Elzen et al., 2014). To restart translation after stress, these 80S need to be dissociated in order to repopulate the pool of free 40S for initiation. Here, the Dom34 splitting system containing the termination factor homologs Dom34 (Pelota in mammals), Hbs1 and the ABC-type ATPase Rli1 (ABCE1) were shown to dissociate vacant ribosomes in yeast (van den Elzen et al., 2014), and mammals (Pisareva et al., 2011). The first eukaryotic hibernation factor, Stm1p, was found in the crystal structures of otherwise empty yeast 80S ribosomes, which were prepared from cells following 10 minutes of glucose starvation (Ben-Shem *et al.*, 2011). Stm1 and its mammalian homolog SERBP1 are clamping the ribosomal subunits together and, similar to bacterial hibernation factors, bind in the mRNA entry channel as well as A and P site of the 40S thereby occupying sites important for translational activity(Ben-Shem et al., 2011).

The non-essential protein Stm1p was shown to have a protective role in *Saccharomyces cerevisiae* (*S.c.*), supporting recovery of translation after quiescence (Balagopal and Parker, 2011; Van Dyke et al., 2013) and a knockout of Stm1 in yeast suppresses requirement of the Dom34 splitting system to restart translation after glucose starvation (van den Elzen et al., 2014). This indicates that idle 80S lacking Stm1 are less stable *in vivo*. Metazoan homologs of Stm1p were also observed bound to inactive ribosomes from *Drosophila melanogaster* (*D.m.*) and *Homo sapiens* (*H.s*.) indicating a high degree of functional conservation (Anger et al., 2013). Notably, these ribosomes also contain tRNA in the ribosomal E site and the elongation factor eEF2 (Anger et al., 2013). Presence of eEF2 and E site tRNA was later also observed on inactive ribosomes derived from rabbit reticulocyte lysates (Brown et al., 2018). In the same system, another type of inactive ribosomes was found containing the poorly characterized protein interferon-related developmental regulator 2 (IFRD2) and tRNA in a newly defined position, near the E site, termed Z site (Brown et al., 2018). However, formation, release and molecular function of the involved RBFs remain enigmatic for all types of hibernating eukaryotic ribosomes.

Recently, another eukaryotic RBF responsible for protecting and recovering translation was discovered: late-annotated short open reading frame 2 (Lso2) in *S.c.* which is highly homologous to coiled-coil domain containing short open reading frame 124 (CCDC124) in human cells (Wang et al., 2018). Lso2 constitutively associates with 80S monosomes and binds in proximity to tRNA, and to rRNA helices H43 and H44 based on chemical crosslinking studies (Wang et al., 2018). This interaction between Lso2 and ribosomes apparently facilitates reactivation of translation upon nutrient upshift and exit from stationary phase. Lack of Lso2 appears to affect translation at the stage of initiation based on a global 4 to 5-fold reduction in ribosome association with most mRNAs in *LSO2* knockout cells (*lso2Δ*) during recovery. Ribosomes from *lso2Δ* show additional functional defects including altered sensitivity to RNase I and altered A site accommodation, as determined by increased incidence of pausing at start codons and enrichment of 21mers in ribosomal profiling – indicative of empty A sites (Wang et al., 2018; Wu et al., 2019). The mode of ribosome interaction and thus the molecular basis for these effects of Lso2 are not understood.

Here, we have structurally characterized the interaction between Lso2 and CCDC124 with eukaryotic 80S ribosomes by single particle cryo-EM to illuminate how binding of these proteins can modulate translational activity during and after starvation. We reconstituted Lso2 with idle 80S ribosomes from purified components and also characterized native idle 80S ribosomes obtained from yeast grown under nutrient-limiting conditions in minimal medium, and from human (HEK293T) cells harvested at high confluency. High resolution structures of the yeast Lso2-80S and the human CCDC124-80S complexes reveal near identical binding modes of Lso2 and CCDC124 to ribosomes. Within idle ribosomes, these factors occupy the P site position of the 40S subunit, including mRNA and tRNAs binding sites. Moreover, they bind the 60S subunit in the A and P sites and reach close to the stalk base and the GTPase activating center (GAC) (Spahn et al., 2004). Surprisingly, we find that a majority of human 80S are occupied with EBP1 – a homolog of the ribosome biogenesis factor Arx1 – bound to the tunnel exit of the 60S (Bradatsch et al., 2007). Notably, we observe in addition to the class containing non-rotated CCDC124-bound 80S, also the previously described idle 80S bound to SERBP1 and eEF2 in the rotated state. These human SERBP1/eEF2 80S are similar to the inactive Stm1-bound ones in the crystal structures of yeast ribosomes (Ben-Shem, 2011), suggesting that in eukaryotes two functionally different pools of idle ribosome exist. Exploring their functional distinction, we show that only Lso2-containing, but not Stm1-containing idle 80S can be readily split by the Dom34 splitting system. This strongly suggests a function of Lso2 in providing a pool of easily recyclable 80S for quick resumption of translation after nutrient starvation.

## Results

### Identification of Lso2 bound to idle yeast 80S ribosomes

For structural analysis of Lso2-bound ribosomes we reconstituted idle 80S ribosomes from *S.c. in vitro* with a 10 × molar excess of purified recombinant Lso2 under defined conditions (see Materials and Methods) and performed single particle cryo-EM. From 3D classification (Supplemental Figure 1A) we obtained one class displaying a non-ribosomal helix-shaped extra density between the large and small subunit (Figure 1A). After refinement, we obtained a structure with 3.4 Å average resolution and local resolution for Lso2 ranging from 3.2 – 4.5 Å (Figure 1A, left, and Supplemental Figure 1D) which allowed us to build an atomic model for Lso2 (Figure 1A, middle, Supplemental Figure 2, Supplemental Table 1).

**Figure 1:**
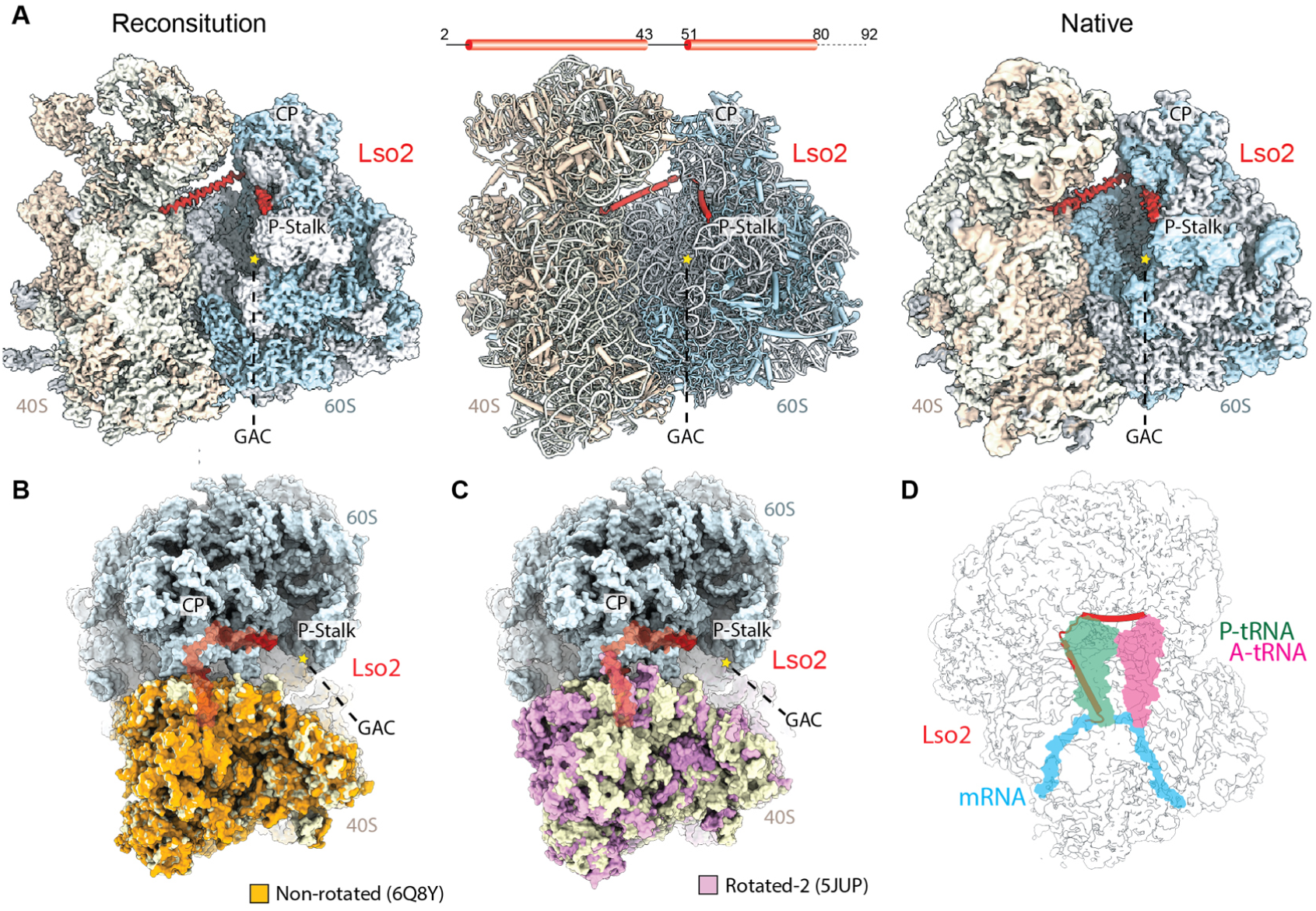
Identification of Lso2 bound to idle 80S ribosomes. A) Left: 3.4 Å resolution cryo-EM map of the *in vitro* reconstituted yeast Lso2-80S complex; α-helical extra density for Lso2 (red) is found in the intersubunit space. Middle: Atomic model of the Lso2-80S complex. Right: 3.5 Å resolution cryo-EM map obtained from a native 80S ribosome preparation from cells grown under nutrient-limiting condition. B-D): Top views of the Lso2-80S structure superimposed with the structure of a yeast 80S in the non-rotated state (B; PDB 6Q8Y (Tesina et al., 2019); C The models are shown in surface representation. C) Top view of Lso2-80S structure superimposed with the structure of a yeast 80S in the rotated-2 state (5JUP,(Abeyrathne et al., 2016), showing differences between SSU rRNA in the two rotational states. All structures were aligned on the 60S subunit. Hallmark features of the 80S ribosome are labelled D) Same view as C, illustrating the position of accommodated A- and P site tRNAs and mRNA (from 5MC6, (Schmidt, 2016)). (CP: central protuberance; GAC: GTPase activating center)

Lso2 is exclusively bound to the inactive 80S in the un-ratcheted/non-rotated conformation as for example observed in 80S ribosomes stalled with an empty A site (Tesina et al., 2019) (Figure1B). This is unusual since idle yeast 80S have normally been observed primarily in rotated states (Figure 1C, independent of the presence of Stm1 (Beckmann et al., 1997; Ben-Shem et al., 2011; Jenner et al., 2012). In our structure, Lso2 comprises two α-helices connected by a short loop (Supplemental Figures 1 and 2). The first Lso2 helix is located in the intersubunit space between small and large subunit. More specifically, it occupies the peptidyl-tRNA binding site (P site) and the mRNA channel in the P and E sites of the small subunit. (Figure 1D). From there, Lso2 bridges over towards the A site of the 60S subunit, whereby the second helix stretches below the central protuberance and extends towards the GAC and the stalk base. In this position, Lso2 would overlap with accommodating A site tRNA (Figure 1D). In conclusion, Lso2 occupies mRNA and tRNA binding sites on the ribosome that need to be accessible for translation, thus corroborating Lso2’s function as a ribosome hibernation factor. Confirming the reconstituted Lso2 80S complex structure, we determined the native structure of Lso2-bound 80S ribosomes (Figure 1A, right), which we found in yeast cells cultured in minimal medium. Extensive classification (Supplemental Figure 1B) yielded a Lso2-containing class at 3.5 Å resolution which was essentially indistinguishable from our *in vitro* reconstituted structure (Supplemental Figure 2, Supplemental Table 1). We conclude that also *in vivo* Lso2 preferably binds empty unrotated 80S ribosomes in the same conformation found *in vitro*.

**Figure 2:**
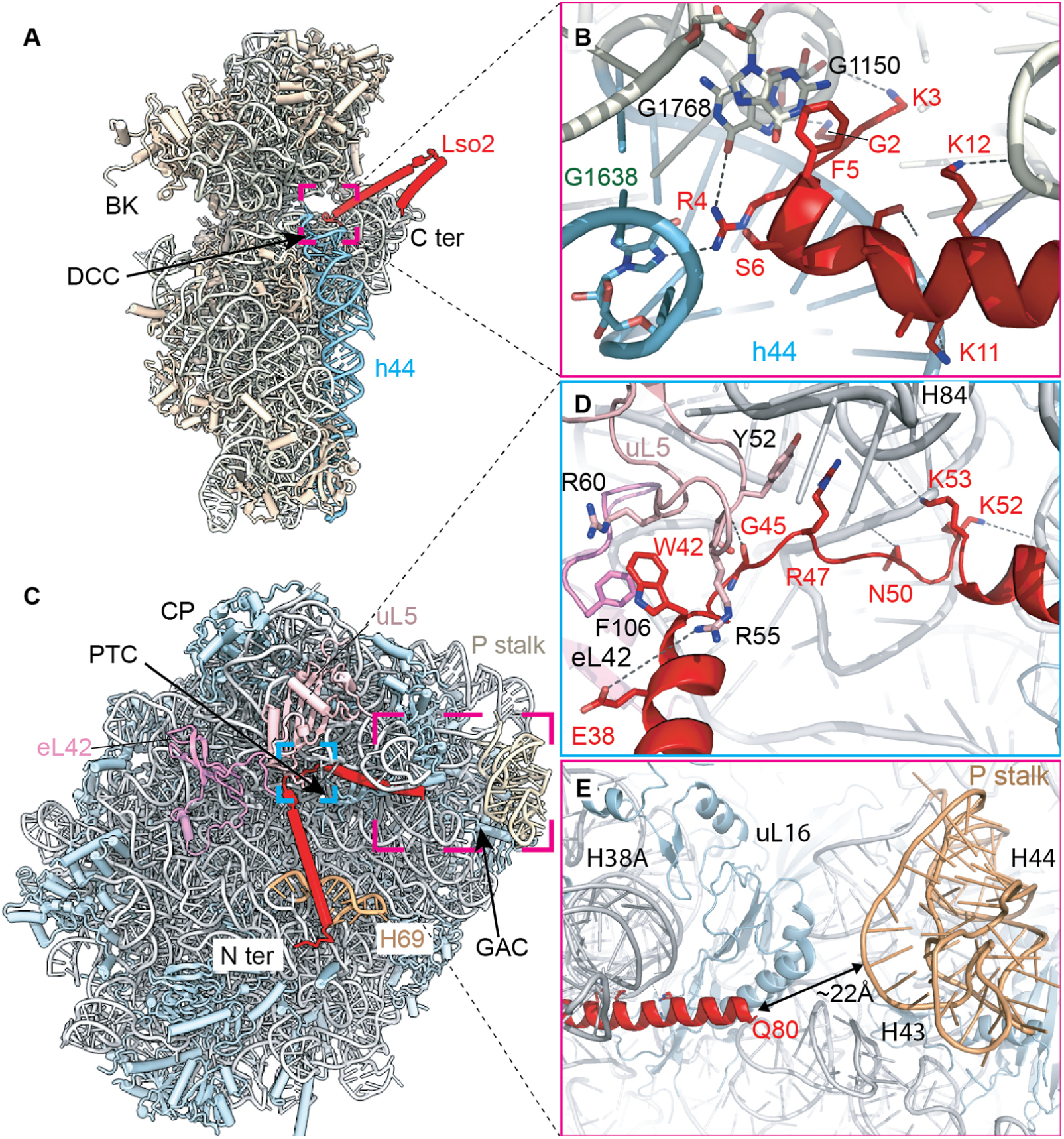
Lso2 interactions with the ribosome. A) Positioning of Lso2 with respect to the small subunit. B) Zoom on interactions of Lso2 in the P- and E sites of the 40S: The Lso2 N-terminus reaches into the cleft between rRNA helices h24, h28 and h44. All the hydrogen bond and salt bridge interactions were indicated with a dashed line; C) Positioning of Lso2 with respect to the large subunit; D) Lso2 binds to the large subunit near the central protuberance (CP) via the helix connecting loop. Interactions involve ribosomal proteins uL5 and eL42 as well as H84 and H85 of 25S rRNA. E) The C-terminal α-helix of Lso2 passes by the major groove of H38A and uL16 towards the stalk base formed by H43 and H44 (GAC) and the P-stalk. The ultimate C terminus of Lso2 is missing, but might be located very close to the P stalk. (BK: beak, DCC: Decoding center, PTC: peptidyl transferase center, GAC: GTPase activating center)

### Lso2 interacts with ribosomal tRNA and mRNA binding sites

The local resolution in the inter-subunit space allowed us unequivocally identify Lso2 and describe its ribosome interactions based on side-chain resolution (Figure 2, Supplemental Figure 2). The main interaction of Lso2 with the 40S subunit is formed by the positively charged Lso2 N-terminus that reaches deeply into the P and E site positions of mRNA (Figure 2A). Here, Lso2 residues G2 to S6 interact in the cleft between rRNA helices h24, h28 and h44 (Figure 2A). In detail, Lso2 F5 is stacking with the bases G1150 of h28 and G1768 of h44; an interaction likely supported by several salt bridges and hydrogen bonds formed between highly enriched positively charged Lso2 residues (K3, R4, K11, K12) and the negatively charged 18S rRNA (Figure 2B). The Lso2 N-terminus thus occupies the site where in an active ribosome, the last two mRNA bases from the E site codon and the first mRNA base of the P site codon would be located.

Protruding from the 40S P site, the density for first Lso2 helix ends near the central protuberance of the large subunit (Figure 2C). The connecting loop constitutes an important interface involving ribosomal proteins uL5 and eL42 as well as 25S rRNA. In detail, Lso2 W42 is accommodated in a pocket formed by R55 and R60 from uL5 and F106 from the C-terminus of eL42 (Figure 2D). Further interactions between Lso2 residues E38, G45 and R47 with uL5 Y52 and R55 as well as between Lso2 N50, K52 and K53 and the phosphate backbone of 25S rRNA helices H84 and H85 are well resolved (Supplemental Figure 2C, 2D). The second straight helix of Lso2 continues along the major groove of H38A (also known as the A site finger) and uL16 towards the stalk base and the P-stalk (Figure 2E). Notably, both the A site finger and uL16 are contact sites for the elbow region of A site tRNA during accommodation and translocation (Frank et al., 2007; Petrov et al., 2008; Whitford et al., 2010). In our structure, these contact sites are blocked by Lso2. RNA crosslinking (ePAR-CLIP) data suggested direct interaction of Lso2 with H43/H44 of the GAC (Wang et al., 2018). This interaction is most likely established by the ultimate C-terminus of Lso2, which is not resolved in our maps due to a high degree of flexibility.

### Two distinct, inactive ribosomal species are present eukaryotes

Since Lso2 is conserved in higher eukaryotes (Wang et al., 2018), we expanded our studies on its role as a starvation factor in the human system. To that end, human 80S ribosomes were prepared from a HEK293T culture with high cell density. Mass spectrometry confirmed the presence of Lso2 homolog CCDC124 as well as SERBP1, eEF2 and EBP1. CCDC124 and EBP1 were also detected in Western Blot analysis (Supplemental Figure 3).

Cryo-EM analysis of this sample revealed two major classes of hibernating 80S ribosomes (Supplemental Figure 4A). One class showed helical density very similar to the density for Lso2 on the yeast ribosome, and the ribosome again adopted the non-rotated state. As expected, the density corresponds to CCDC124. Interestingly, the same class also displayed density corresponding to EBP1 at the peptide exit site (Barrio-Garcia et al., 2016; Bradatsch et al., 2012; Greber et al., 2016), as well as tRNA in the E site. The second class also contained EBP1 and E site tRNA, but instead of CCDC124, SERBP1 and translation elongation factor eEF2 were present and ribosomes were stabilized in the rotated state, as previously observed in *D. m*. and *H. s*. inactive 80S ribosome cryo-EM maps (Anger et al., 2013; Brown et al., 2018).

Both classes, as well as a merged class of all EBP1 containing particles, were refined independently (Supplemental Figure 4B, 4C, 4D) and a molecular model was built for the human CCDC124-EBP-80S, a SERBP1-eEF2-EBP1-80S complex as well as an EBP1-80S complex for which all EBP-containing particles were refined (Figure 3A, 3B, Supplemental Figure 5 and Supplemental Table 1).

**Figure 3:**
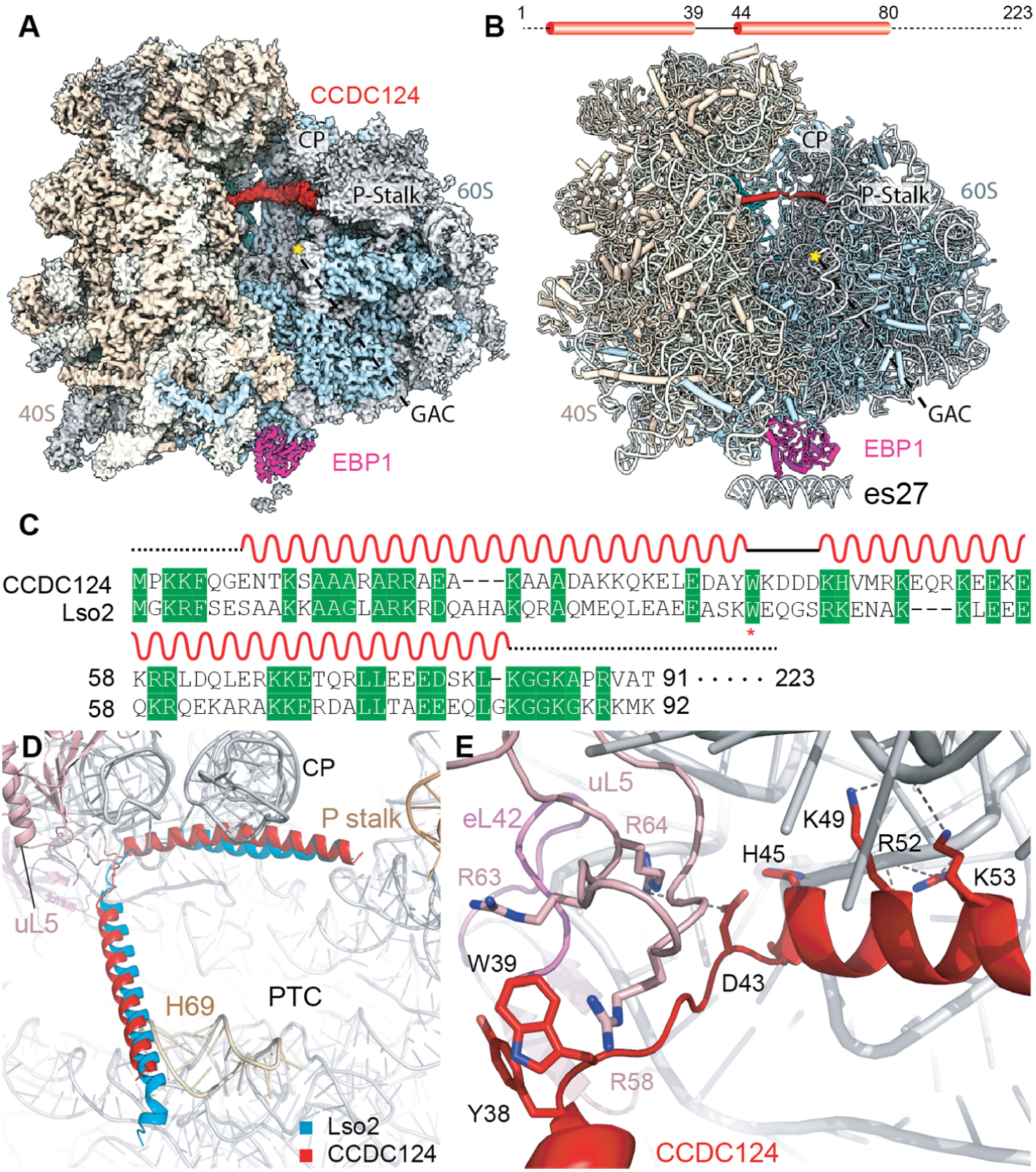
Ribosome binding of Lso2 and CCDC124 is evolutionary conserved. A) 3.0 Å resolution cryo-EM map of the human CCDC124-EBP1-80S ribosome. The reconstruction also contains E site tRNA. The CCDC124 density is displayed at a different contour level than the ribosome map. B) Atomic model of the CCDC124-EBP1-80S complex. C) Conserved residues shown in a sequence alignment of CCDC124 with Lso2. D) Superposition of Lso2-80S with CCDC124-80S models based on 25S rRNA shows a very similar positioning indicating a highly conserved binding mode. CP = central protuberance; PTC = Peptidyl transferase center.

### The ribosome binding mode of Lso2 and CCDC124 is highly conserved

As for the yeast Lso2-80S complex, the local resolution was sufficient to assign the helical density unambiguously to CCDC124 and describe its ribosome interactions (Figure 3E, Supplemental Figures 4B-D and 5, Supplemental Table 1). Position and orientation of CCDC124 in the A and P sites of the 40S was near identical compared to the yeast homolog, indicating a conserved binding mode (Figure 3). Ribosome binding is mediated by residues conserved from *S.c.* to *H.s.* (Figure 3C), including W39 (W42 in *S.c*.) which plays a key role in establishing the main contact to the 60S in the P site.

In detail, the CCDC124 N-terminal helix is occupying a similar space as Lso2 in the P and E site mRNA position (Figure 3D), yet the electron density for the N-terminal helix was weaker than for the C-terminal one, most likely due to higher overall flexibility of the 40S subunit. The resolution of 3.0 Å was sufficient for molecular model building starting from K11. As with Lso2, the first helix stretches over towards the 60S subunit. At the connecting loop, W39 of CCDC124 (similar to W42 in Lso2) is accommodated on the 60S in a binding pocket formed by uL5 R58 and R63 (Supplemental Figure 5C). Unlike the yeast Lso2 contacts, however, eL42 is not involved in CCDC124 binding (Figure 3E). The stacking interaction between W42 of Lso2 and F106 of eL42 is accordingly not conserved in the human complex. Yet, it is compensated for by stacking between Y38 and W39 of CCDC124. In general, the helix-connecting loop adopts a slightly different path, though the second helix is observed in a nearly identical position as in Lso2 (Figure 3D). Analogous to what we observe in the Lso2 structures, presence of CCDC124 excludes A site tRNA accommodation, and again, the extended C-terminal tail is not resolved (Figure 3D, Supplemental Figure 5D). In conclusion, CCDC124 occupies the binding interfaces of A and P site tRNA, as well as mRNA on the human ribosome concurrent with the Lso2 binding site in yeast (Figure 3D and 1D).

The second major class of hibernating human ribosomes we observed and refined to 3.1 Å was in agreement with previous reports in the rotated state, more specifically the “rotated-2 state” as occurring when the 80S is occupied with hybrid A/P and P/E tRNAs (Figure 1C) (Abeyrathne et al., 2016). In addition to a type-II tRNA displaying an extended V-loop in the E site, it contains eEF2 and SERBP1 (Figure 4). Like Stm1, SERBP1 together with eEF2 binds in the mRNA entry channel and prevents mRNA binding in the A and P sites (Anger et al., 2013; Balagopal and Parker, 2011; Ben-Shem et al., 2011; Brown et al., 2018; Van Dyke et al., 2013). Importantly, binding of SERBP1 and CCDC124 is mutually exclusive.

**Figure 4:**
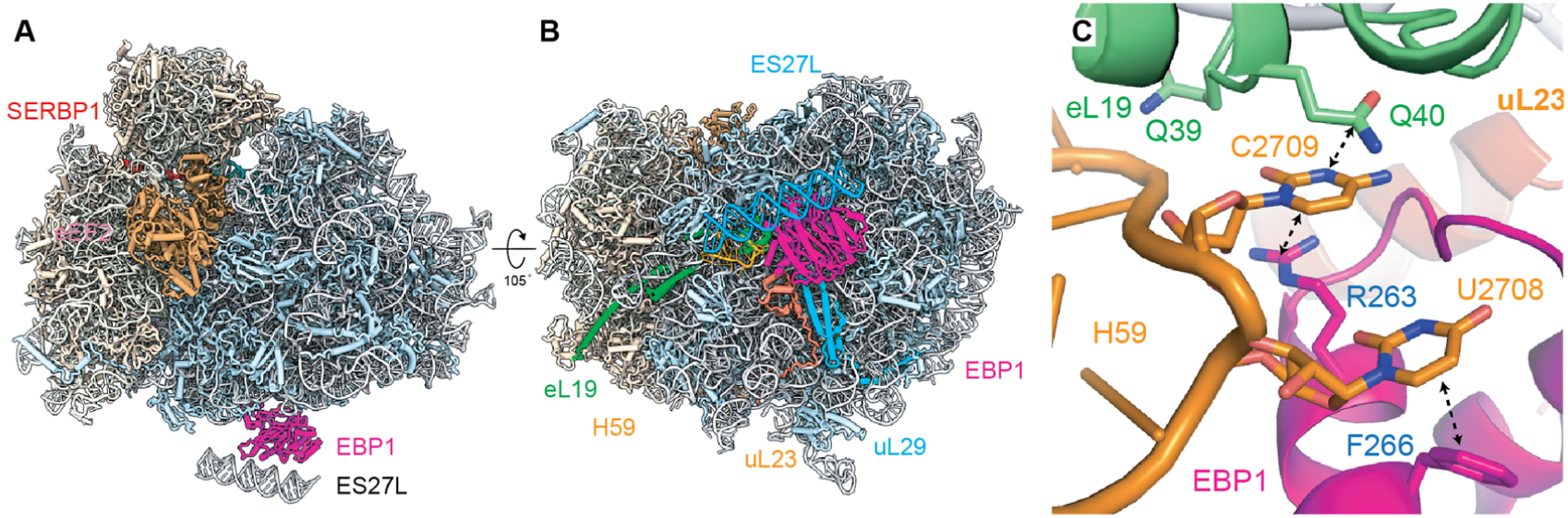
Hibernating human ribosomes are bound to EBP1 on the peptide exit site. A) side and bottom views of the human eEF2-SERBP1-EBP1-80S model, built from a 2.9 Å resolution cryo-EM map. B) EBP1 coordinates a part of rRNA expansion segment ES27L below the peptide tunnel exit. A dummy RNA helix was used as a placeholder for ES27L. C) Zoom on the EBP1 insertion domain reaching into a cleft between uL23 and H59.

### EBP1 binds to the peptide exit site of hibernating human ribosomes

We note that under our conditions the human ribosomes happen to be stably associated with EBP1 at the tunnel exit (Figure 4, Supplemental Figure 6), independent of ribosomal state and factors occupying mRNA and tRNA binding sites. EBP1 is the human homologue of the yeast ribosome biogenesis factor Arx1 (Bradatsch et al., 2012; Kowalinski et al., 2007a) which also binds to the ribosomal exit site and is related to methionine aminopeptidases (MetAPs)(Kowalinski et al., 2007a; Kowalinski et al., 2007b). In addition, EBP1 has been described to be involved in cell proliferation (Squatrito et al., 2004) and human cancer (Nguyen et al., 2018).

We obtained a structure of EBP1-80S at 2.9 Å overall resolution, showing that EBP1 is anchored to the peptide exit site through ribosomal proteins uL23, eL19, uL29 as well as 28S rRNA helix H59 (Figure 4C, Supplemental Figure 6). Also similar to Arx1 in yeast, it coordinates a part of the flexible rRNA expansion segment ES27L in a defined position below the tunnel exit. Its previously described insertion domain (residues 250-305) (Kowalinski et al., 2007a) interacts with rRNA helix H59 in a rearranged conformation, and reaches into a cleft between uL23 and H59 (Supplemental Figure 6A). On the tip of H59 the base U2708 is flipped out and stacks with EBP1 F266 (Figure 4C) while C2709 is sandwiched between R263 of EBP1 and Q40 of eL19. These interactions are supported by additional contacts between its MetAP-like domain and uL23 and uL29 (Figure 4B). Interestingly, EBP1 shares a similar interaction mode with the ribosome as nascent chain interacting factors such as RAC, SRP, Sec61 or the NatA complex (Becker et al., 2009; Knorr et al., 2019; Voorhees and Hegde, 2016; Zhang et al., 2014), and the presence of EBP1 would not allow simultaneous binding of any of the above mentioned factors (Supplemental Figure 6C).

### Lso2-bound but not Stm1-bound 80S are split by canonical recycling factors

Our cryo-EM analysis of human hibernating 80S suggests that in eukaryotes, at least two clearly distinguishable populations of idle, translationally repressed 80S exist. 80S bound to Stm1 and SERBP1/eEF2, and 80S bound to Lso2 (CCDC124). Besides the different factor compositions, their main difference is the conformation of the ribosome: Stm1/SERBP1-containing 80S so far have been exclusively observed in rotated states (Abeyrathne et al., 2016; Anger et al., 2013; Ben-Shem et al., 2011; Brown et al., 2018), whereas the Lso2- or CCDC124-bound 80S are in the non-rotated state (Figure 1B, 1C), similar to the POST-translocational state with tRNAs in P and E sites and an empty A site (Budkevich et al., 2014; Tesina et al., 2019). Thereby, Lso2 (CCDC124) on 80S occupy a position that would on one hand exclude simultaneous presence of the basic translation machinery (A site tRNA, P site tRNA, mRNA) (Figure 1D), on the other hand stabilize the non-rotated ribosome conformation, which is required for binding of Dom34 splitting system (Figure 5A and 5B).

**Figure 5:**
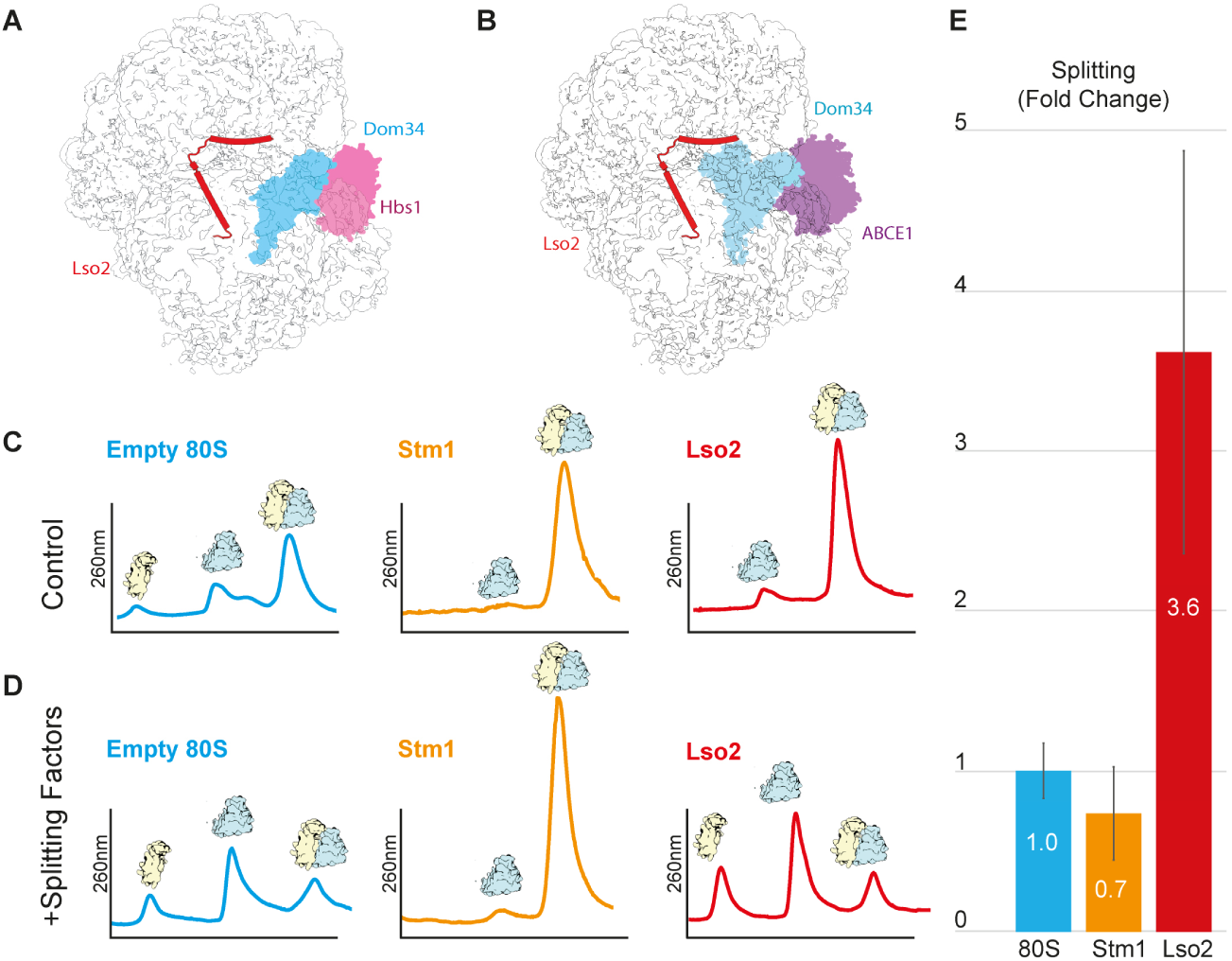
*In vitro* splitting assay with Lso2-80S and Stm1-80S. A and B): The Lso2-80S in non-rotated state (displayed as cartoon) was overlaid with ribosome rescue factors Dom34 and Hbs1 (Becker et al., 2011) or Dom34 and ABCE1 (B; PDB: 3J16; (Becker et al., 2012). In both conformations, Dom34 could accommodate within the A site of the 40S and would not clash with Lso2, leading to the hypothesis that the Dom34 splitting system preferably splits Lso2-80S. C and D): representative sucrose gradient profiles from splitting assays carried out without (C) or with splitting factors (D). E) Quantification of relative splitting, displayed in fold change, normalized to propensity of splitting observed in the control (puromycin-treated) empty 80S (see also Supplemental Figure 7 and Supplemental Figure 8). All experiments were performed in triplicates.

We tested our hypothesis, that the Lso2-bound 80S ribosome is the preferred substrate for the Dom34 splitting system, using an *in vitro* ribosome splitting assay. To that end, 80S ribosomes were incubated with a 5-fold molar excess of purified Dom34, Hbs1-GTP, ABCE1-ATP and, to prevent subunit re-association, eIF6. Splitting reactions were subjected to sucrose density gradient centrifugation and UV profiles of the gradients were generated at 254 nm (Supplemental Figure 7).

Splitting assays were performed with reconstituted Lso2-80S ribosomes, Stm1-enriched 80S (Ben-Shem et al., 2011) and puromycin/high salt treated empty 80S ribosomes as a control. Using near-physiological splitting conditions, control 80S were partially split, whereas almost quantitative splitting of Lso2-80S was observed (Figure 5D and 5E, and Supplemental Figure 8). Conversely, Stm1-80S complexes were found to be essentially resistant to splitting by the Dom34 system (Figure 5D and 5E, and Supplemental Figure 8). To confirm that the non-splittable fraction in the Stm1-80S population is indeed containing Stm1 bridging the 40S and 60S, we performed cryo-EM on the Stm1-80S from splitting assays with and without splitting factors. In both cases, we observe that a vast majority (>90%) of splitting-resistant 80S adopted the same “rotated-2 state” as the human eEF2 and SERBP1 containing ribosomes. Here, in more than 50 % of these ribosomes Stm1 could be observed in the subunit-bridging conformation crossing the mRNA channel (Supplemental Figure 9). In conclusion, these results strongly suggest that the Lso2 or CCDC124-bound ribosomes, which are stabilized in the non-rotated state, are indeed preferred substrates for the Dom34 splitting system.

## Discussion

While for bacteria numerous hibernation factors have been characterized (for review see e.g. (Prossliner et al., 2018), knowledge about eukaryotic equivalents is so far limited to Stm1 in yeast and SERBP1/eEF2 as well as the recently discovered IFRD2/Z-site tRNA (Ben-Shem et al., 2011; Brown et al., 2018) in mammals. In analogy to bacterial counterparts, both SERBP1/eEF2 and IFRD2/Z-tRNA occupy crucial active sites on the ribosome (Brown et al., 2018). While structural data strongly suggested that hibernation factors act by maintaining ribosomes in a translation-incompetent state, the mechanisms of recovering translation-competent ribosomes have remained enigmatic.

In this work, we present Lso2 and CCDC124 in yeast and human cells as new eukaryotic-specific ribosome hibernation factors that play an active role in translation recovery (Wang et al., 2018). Our cryo-EM structures show, that Lso2 and CCDC124 occupy the binding sites of mRNA and tRNA, thereby resembling the mode of action of known bacterial hibernation factors. However, while RMF, HPF, RaiA or the IHPF N-terminal domain all target the mRNA path and decoding center of the small subunit, Lso2 and CCDC124 bind to both, 40S and 60S subunits and exclusively stabilize the non-rotated conformation of the 80S ribosome.

Independent of the ribosomal state and binding of hibernation factors SERBP-1 or CCDC124, we observe EBP1 bound to the peptide tunnel exit of hibernating human ribosomes. The binding mode we observe is similar to that of other exit site binding proteins which interact with the nascent chain. As a result, this binding mode excludes interaction of the majority of co-translational acting factors to the ribosome, consistent with a role as hibernation factor that prevents unproductive sequestration of these factors to idle ribosomes. Another potential role of EBP1 may be in preventing ubiquitination of the ribosomal protein uL29 of the inactive ribosome, thereby contributing to the coordination of its propensity for ribophagy (Brandman and Hegde, 2016). Yet, the exact circumstances and mechanism of how EBP1 is recruited to a dormant ribosome and what triggers its dissociation before or during re-entering the translation cycle remains to be elucidated.

We reasoned that keeping dormant ribosomes in the non-rotated state may be a prerequisite for the preferred reactivation by the Dom34 splitting system. Indeed, using *in vitro* splitting assays we could show that Lso2-bound 80S are highly susceptible to recycling. In contrast, 80S enriched in Stm1 – as occurring after short glucose starvation (Ben-Shem et al., 2011)– could not be split, which is rationalized by our structures of Stm1-80S resembling a distinct rotated-2 state (Abeyrathne et al., 2016). This is highly consistent from a structural perspective, since Dom34-Hbs1 and Dom34-ABCE1 complexes preferably bind to ribosomes in the non-rotated POST state (Becker et al., 2011; Becker et al., 2012; Brown et al., 2015b; Hilal et al., 2016; Shao et al., 2016) but clash with the rotated state that is stabilized by Stm1 (Supplemental Figure 10). Moreover, presence of Stm1 or SERBP1 to a large extend results in stable eEF2 binding in the translation factor binding site in mammals (Anger et al., 2013; Brown et al., 2018) and, at least *in vitro*, also in yeast (Hayashi *et al.*, 2018). This would further block access of Dom34 and render these ribosomes resistant to splitting by this system.

These findings are also in good agreement with the initial observation that absence of Lso2 delays recovery of translation after starvation (Wang et al., 2018). Moreover, Lso2 depleted cells accumulate idle 80S ribosomes and are decreased 5-fold in global initiation as analyzed by ribosome profiling (Wang et al., 2018). Intriguingly, a study demonstrated that a deletion of Dom34 has similar effects on recovery from glucose-depletion in yeast as deletion of Lso2, and that idle ribosomes isolated from glucose-starved yeast were substrates for splitting by the Dom34 system (van den Elzen *et al.*, 2014). Moreover, deletion of Stm1 suppresses the requirement for Dom34 and Hbs1 in translation recovery, indicating that presence of Stm1 in ribosomes antagonizes the rapid splitting action of Dom34 and Hbs1. In support of this antagonizing role, overexpression of Stm1 in a *dom34Δ* strain leads to a growth defect (Balagopal and Parker, 2011), likely due to an accumulation of Stm1-80S and the inability to split the remaining pool of 80S by the Dom34 system. In combination with these data, our findings strongly suggest that Stm1-80S represent a different pool of dormant ribosomes that is most likely not readily split by the Dom34 system. Taken together, we conclude that Lso2 acts in protecting inactive 80S ribosomes as a hibernation factor, yet in contrast to Stm1, Lso2 stabilizes them in a rapid to recycle state for timely re-entrance into the active translation cycle. The question remains, what the actual signals are for both, Stm1 and Lso2, to engage with idle 80S in their active subunit-clamping conformations. Both proteins are likely to be constitutively associated with ribosomes, even during the translation cycle. Stm1 for example was shown to repress translation, in yeast perhaps by modulating access of translation elongation factor eEF3 (Hayashi et al., 2018; Van Dyke et al., 2013), and to also play a role in mRNA deadenylation (Balagopal and Parker, 2011). A role in early elongation was also attributed to Lso2 (Wang et al., 2018). Apparently, when already bound to 80S ribosomes, both factors can quickly generate a pool of protected dormant 80S that is either inaccessible (Stm1-80S) or accessible (Lso2-80S) to the Dom34 splitting system. Yet signals decisive to generation and disassembly of these seemingly redundant, albeit alternate inactive 80S populations, for example relative population sizes or specific stress conditions are unclear. Furthermore, Lso2-80S must be protected from recurrent recycling to avoid repeated splitting and formation of these 80S. Whether this is regulated by the energy levels in the cell or by other specific signals should be the subject of further investigation.

## Materials and Methods

### Purification of recombinant Lso2

Lso2 was overexpressed in *E. coli* BL21 (DE3) grown in LB media supplemented with ampicillin. Cells were induced at an OD_600_ of 0.05 and grown at 37°C to mid log phase. Protein expression was induced by the addition of IPTG. Cells were harvested after two hours of protein expression by centrifugation at 3,500 × *G*. Cell pellets were re-suspended in lysis buffer (50 mM Tris-HCl pH 8.0, 300 mM NaCl, 2 mM β-mercaptoethanol (β-Me)) and lysed using Microfluidics M-110L microfluidizer. The membrane fraction was removed by centrifugation at 34,000 × *G* for 45 min. Clarified lysates were then loaded onto nickel resin equilibrated with five column volumes (CV) of wash buffer (30mM Tris-HCl pH 8.0, 300 mM NaCl, 20 mM Imidizole, 2 mM β-Me). Lso2 was cleaved in batch mode by addition of de-ubiquitin protease over night at 4°C and eluted in wash buffer lacking imidazole. The eluate was concentrated to 1 mL using a Millipore 3 MWCO concentrator and applied on a SuperDex 75 gelfiltration colum in 20 mM Tris-HCl pH 8.0, 150 mM NaCl, 2 mM β-Me, yielding 1.1 mg purified Lso2 from 0.5 L culture at a concentration of 3 mg/ml.

### Purification of ribosomal subunits

In short, clarified cytoplasmic lysates obtained from isogenic *S. cerevisiae* S288C cells (MATα; HIS3; LEU2; ura3-52; TRP1; GAL2) were spun through a sucrose cushion (1M sucrose, 30 mM Tris-HCl pH 7.0, 500 mM KOAc, 25 mM Mg(OAc)_2_, 5mM β-Me, 0.1% Nikkol, 10 µg/µL Cycloheximide) at 290,000 × *G* for 45 min. The ribosomal pellet was re-suspended in buffer (50 mM Tris-HCl pH 7.4, 500 mM KOAc, 2 mM MgCl_2_, 2 mM DTT) and treated with puromycin (1 mM final concentration) for 15 min on ice, and 10 min at 37°C. Samples were then loaded onto 10-40% sucrose gradients (50 mM HEPES pH 7.4, 500 mM KOAc, 5 mM MgCl_2_, 0.1 mM EGTA, 2 mM DTT) and centrifuged for 3 h at 284,000 × *G*. Gradient fractionation was carried out and 40/60S subunits were concentrated using Amicon 100k MWCO concentrators against grid buffer (20 mM HEPES 7.4, 100 mM KOAc, 2.5 mM Mg(OAc)_2_, 250 mM Sucrose, 2 mM DTT).

### Reconstitution of the Lso2-80S complex

Purified ribosomal subunits were mixed at 1:1 molar ratios and incubated under re-association conditions in grid buffer containing 10 mM Mg(OAc)_2_, 0.1 % Nikkol for 10 min. Afterwards a ten-fold molar excess of purified Lso2 was added 10 min prior to blotting.

### Native Lso2-80S complexes from S. cerevisiae

We identified Lso2-containing ribosomes by cryo-EM in several samples, where ribosomal complexes were purified from yeast cells grown in minimal medium and overexpressing different target proteins on plasmids. The sample analyzed here was initially targeted at obtaining Upf1-containing ribosomal complexes from BY4741 yeast cells (*MATα; ura3Δ0; leu2Δ0; his3Δ1; met15Δ0; YMR080c::kanMX4*) harboring the plasmids pKB510 (Serdar et al., 2016) overexpressing a nonsense-mediated mRNA decay reporter and pKB607 overexpressing a FLAG-tagged ATP-hydrolysis-deficient mutant. Cryo-EM analysis of the elution fractions revealed a vast majority of idle 80S ribosomes lacking any density for Upf1, however, revealing a subclass of Lso2-bound ribosomes.

In detail, cells were grown in minimal medium (Yeast Nitrogen Base; -Leu-Ura dropout medium and 2 % glucose) at 30 °C to an OD_600_ of about 0.75. Lysis was performed with Microfluidics M-110L microfluidizer using lysis buffer (20 mM HEPES pH 7.4, 100 mM KOAc, 10 mM MgCl_2_, 0.5% Triton, 1:1000 protease inhibitor pill (Roche: 04-693-132-001). For the preparation 40 g of lysed cell powder was used and a cytosolic S100 fraction was prepared. First, membrane fractions were removed by centrifugation at 28714 × *G* for 15 min, then cytosolic fractions were clarified by centrifugation at 92387 × *G* for 20 min. The “S100” was added on 300 µl of magnetic FLAG beads (Sigma-M8823) equilibrated with lysis buffer and incubated for 2 h at 4 °C. After washing three times with lysis buffer lacking Triton X-100, the sample was eluted with FLAG peptide (Sigma F4799). Ribosomal eluate were spun through a 750 mM sucrose cushion prepared in elution buffer for 45’ at 290,000 × *G*. Pellets were resuspended in elution buffer and adjusted to a final concentration of ∼*4 A*_260_/ml for cryo-EM sample preparation.

As mentioned above, only Lso2 could be visualized in this sample as additional ribosome binder. Similar observations were made when using the same protocol to obtain ribosomal complexes with other tagged proteins indicating that the presence of Lso2 on vacant ribosomes is independent of the nature of the tagged bait protein.

### Native human hibernation complexes

5 × 15cm plates of HEK293 Flp-In T-REx cells (ThermoFisher Scientific, R78007) at 80% confluency were harvested in 15mL DMEM media by trypsinization, and pelleted by centrifugation for 10 min at 150 × *G*. Cells were washed with ice cold PBS and pelleted again before re-suspending in 0.75mL lysis buffer (20 mM HEPES/NaOH pH 7.4, 100 mM KOAc, 10 mM Mg(OAc)_2_, 125 mM Sucrose, 1 mM DTT, 0.5 mM PMSF, 0.5 % IGEPAL, protease inhibitor tablet). Cells were incubated for 30 min at 4 °C before pelleting the cell debris for 15 min at 21,000 × *G*. The clarified lysate was distributed on top of 10-50% sucrose gradients prepared with lysis buffer lacking IGEPAL and centrifuged for 3 hours at 284,000 × *G*. 80S fractions were collected, combined, and pelleted through a 1M sucrose cushion of lysis buffer lacking IGEPAL by centrifugation for 1 hour at 540,000 × *G*. Pellets were re-suspended in 20 µL lysis buffer lacking IGEPAL at a final concentration of 130 *A*_260 nm_ ml^−1^. For cryo grids the concentration was adjusted to 4 A_260_/ml.

### Purification of idle 80S ribosomes with Stm1

Wild type BY4741 *S. cerivisiae* cells were prepared exactly as described previously (Ben-Shem et al., 2011). In short, cells were grown to mid-log phase in YPD before pelleting at 30°C and incubating in YP for a further 10 min at 30°C. Cells were pelleted and washed three times in wash buffer (30mM HEPES pH 7.4, 50mM KCl, 2.5mM Mg(OAc)_2_, 0.5mM EDTA, 2mM DTT, protease inhibitor tablet, 10% Glycerol.) After washing, cells were dropped in liquid nitrogen and lysed using FREEZER MILL SPECS and stored at −80°C. Clarified lysates re-suspended in wash buffer were loaded onto 10-50% Sucrose gradients in wash buffer lacking glycerol. 80S peaks were pelleted through a 1M sucrose cushion prepared in wash buffer, by centrifugation at 417,000 × *G* for 45 minutes. Finally, purified ribosomal pellets were re-suspended in storage buffer (20 mM HEPES pH 7.5, 100 mM KOAc, 5 mM Mg(OAc)_2_, 1 mM DTT). Aliquots were flash frozen and stored at −80°C.

### Purification of Puromycin treated 80S ribosomes

*S. cerivisiae* BY4741 cells were grown to mid log phase in YPD at 30°C, and harvested at a final OD_600_ of 2.5. Cells were pelleted and washed once with water, once with 1% KCl, and re-suspended in 100mM Tris HCl pH 7.95, 10mM DTT and incubated at room temperature for 15 minutes before a final pelleting step. Cells were re-suspended in lysis buffer (10mM HEPES pH 7.5, 100mM KOAc, 7.5mM Mg(OAc)_2_, 125mM Sucrose, 1mM DTT, 0.5mM PMSF, complete EDTA free protease inhibitor tables), before being lysed using Microfluidics M-110L microfluidizer. Lysates were clarified via centrifugation at 4°C, first at 27,000 × *G* for 15 minutes, and again at 150,000 × *G* for 30 minutes. Ribosomal fractions were then isolated by centrifugation through a double layer 1.5M/2M Sucrose cushion (20mM HEPES pH 7.5, 500mM KOAc, 5mM Mg(OAc)_2_,1mM DTT, 0.5mM PMSF) at 246,500 × *G* for 21 hours. Supernatant fractions were discarded and ribosomal pellets were re-suspended in nuclease free water. Ribosomes were mixed 1:1 with 2x Puromycin buffer (40mM HEPES pH7.5, 1M KOAc, 25mM Mg(OAc)_2_, 2mM Puromycin 2mM DTT, and Amicon Anti-RNase (AM2692)) and incubated for 30 minutes at room temperature. Puromycin treated ribosomes were loaded onto 10-40% sucrose gradients in buffer conditions matching previous sucrose cushion, and subjected to ultracentrifugation at 4°C, 21.000 × *G* for 20 hours. 80S fractions were manually harvested from the gradients by monitoring absorption at 260nm, and ribosomes were pelleted at 417,000 × *G* at 4°C for 45 min Finally, puromycin treated 80S pellets were re-suspeded in storage buffer (20 mM HEPES pH 7.5, 100 mM KOAc, 5 mM Mg(OAc)_2_, 1 mM DTT). Aliquots were flash frozen and stored at −80°C.

### Purification of splitting factors

*S. cerevisiae* Dom34 was expressed with a C-terminal His tag in pET21a(+) vector and was purified from Rosetta2(DE3) *E. coli* with Ni-NTA affinity chromatography as previously reported (Becker et al., 2011).

*S. cerevisiae* Hbs1 was expressed with an N-terminal His tag in pET28b(+) vecor and was purified from Rosetta2(DE3) *E. coli* with Ni-NTA affinity chromatography as previously described (Becker et al., 2011). Both proteins were purified in a final buffer of 20 mM Tris pH7.5, 200mM NaCl, 5mM β-ME, 0.1mM PMSF, aliquots were flash frozen and stored at – 80°C.

*S. cerevisiae* Rli1 (ABCE1) in high copy pYES2Rli1WCGα vector was overexpressed in *S. cerivisiae* strain WCGα cells were grown in YP -ura, 2% galactose, 1% raffinose media at 30°C to mid-log phase and were harvested at a final OD_600_ of 1.0. Cell pellets were washed with water before flash freezing and storing at −80°C. For lysis, cell pellets were thawed and washed once with 1% KCl for cell wall destabilization, before resuspending in 100mM Tris pH 8.0, 14mM β-MEand incubating at room temperature for 15 minutes. Subsequently cells were pelleted and re-suspended in Lysis Buffer (75mM HEPES pH 8.0, 300mM NaCl, 5mM β-ME, 1% Tween, 20mM Imidizole, 2mM MgCl_2_, 10%Glycerol) and lysed using Microfluidics M-110L microfluidizer. Lysates were clarified by centrifugation at 25,000 × *G* for 10 minutes and filtered through 0.45µm filter before loading onto HisTrap-HP 5mL affinity column column using the Äkta pure HPCL. The column was washed with wash buffer (50mM HEPES pH 8.0, 500mM NaCl, 5mM β-ME, 20mM Imidizole, 2mM MgCl_2_, 10% glycerol) before eluting over 40 minutes with a 0-100% gradient from wash to elution buffer (wash buffer with 300mM imidazole). Peak fractions were concentrated using Amicon® 50k MWCO concentrator (UFC805024) before loading onto Superdex200 for size exclusion chromatography. Aliquots of pure ABCE1 in 20mM Tris pH 7.5, 200mM NaCl, 5mM β-ME, and 5% glycerol were flash frozen and stored at −80°C.

*S. cerevisiae* eIF6 was cloned into p7XC3GH (addgene#47066) fused at the C-terminus to a 3C protease cleavage site, GFP, and 10-His. The plasmid was transformed into *E. coli* BL21 (DE3), which was grown at 37°C to mid log phase. Temperature was reduced to 16°C and protein overexpression was induced with IPTG for overnight expression. Cells were harvested (4400 × *G*, 4°C, 8 minutes), washed with Phosphate buffered saline, and re-suspended in lysis buffer (20mM Tris pH8.0, 300mM NaCl, 2mM β-Me, and complete EDTA free protease inhibitor tablet) before lysing with Microfluidics M-110L microfluidizer. Lysates were clarified by centrifugation at 30596 × *G* at 4°C for 20 minutes. The supernatant was loaded onto TALON® metal affinity resin equilibrated in lysis buffer and incubated on a rotating wheel at 4°C for 40 minutes. Flow through was collected and resin was washed three times with lysis buffer + 10mM imidazole before incubating with elution buffer (lysis buffer, 10mM imidazole, 2.5mg 3C protease) for 30 minutes at 4°C. Eluted protein was concentrated with Amicon® 10k MWPO concentrator (UFC901096) to 1mL before loading onto Superdex200 for size exclusion chromatography in final buffer (50mM HEPES pH 7.5, 500mM KCl, 2mM MgCl_2_, 2mM β-ME). Aliquots were flash frozen and stored at −80°C.

### Splitting Assay

Splitting Assays were assembled in 50 µL reactions in a final SA buffer (20 mM Tris pH 7.5, 100 mM KOAc, 4 mM Mg(OAc)_2_, 5 mM β-ME, with 5 pmol purified ribosomes (Stm1-bound, Lso2-bound, or Puromycin treated), and a 5 × molar excess of each factor included in the reaction. Compensation buffers were calculated for each reaction to ensure standardization of buffer conditions, regardless of component variability. Control reactions were assembled with ribosomes and eIF6. Splitting reactions included ribosomes eIF6, 10 mM ATP, 10 mM GTP and splitting factors (Dom34, Hbs1 and ABCE1). Reactions were incubated on ice for 30 min before loading onto 10-50% sucrose SA buffer gradients and subjected to ultracentrifugation for 3 h at 284,000 × *G.* Sucrose density gradients were subjected to mechanical fractionation and UV spectroscopy). Quantification of peaks was performed by estimating the integral using the trapezoid rule (Kalambet et al., 2018). Let (x_n_ | A_n_) be the data points recorded with x being the distance along gradient and A the absorption at 260 nm. The area S_ab_ under one peak from x_a_ to x_b_ was approximated as

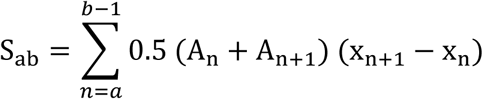

Relative splitting efficiencies were calculated as ratios of peak areas and averaged across experiments. Errors shown in normalized results were estimated assuming linear propagation of statistical uncertainties.

### Cryo-EM analysis

For all yeast samples, ∼ 2.5-6 *A*_260 nm_ ml^−1^ of ribosome were applied to 2 nm precoated Quantifoil R3/3 holey carbon support grids. Data were collected on a Titan Krios TEM (Thermo Fisher) equipped with a Falcon II direct detector at 300 keV under low-dose conditions using approximately 25 electrons per Å^2^ for ten frames in total (Supplemental Table 1). The defocus range was between −1.1 to −2.3 µm and for semi-automated data acquisition the software EM-TOOLS (TVIPS) was used. Magnification settings resulted in a pixel size of 1.084 Å pixel^−1^. Original image stacks were summed and corrected for drift and beam-induced motion at the micrograph level using MotionCorr2 (Zheng et al., 2017). The estimation of contrast transfer function (CTF) and the resolution range of each micrograph was performed with Gctf (Zhang, 2016).

For the human sample, 4 A_260 nm_ ml^−1^ of ribosomes were applied to 2 nm precoated Quantifoil R3/3 holey carbon support grids. Data were collected on a Titan Krios TEM (Thermo Fisher) equipped with a Falcon III direct detector at 300 keV (Supplemental Table 1). The defocus range was between −0.8 to −3.2 µm and for semi-automated data acquisition the software EPU (Thermo Fisher). Frame alignment was performed using MotionCorr2 (Zheng et al., 2017). The estimation of contrast transfer function (CTF) was performed with Gctf (Zhang, 2016).

### Structure of the in vitro reconstituted yeast Lso2-80S complex

After manual screening for ice quality, 10,313 micrographs were used for automated particle picking in Gautomatch (http://www.mrc-lmb.cam.ac.uk/kzhang/) yielding 1,413,783 initial particles. Upon 2D classification in RELION 3.0, 781,257 particles were selected for a consensus 3D refinement. After 3D classification, a Lso2-containing class (88523 particles) was obtained and refined including CTF refinement to an average resolution of 3.4 Å with local resolution ranging from 3-7 Å (3.2-4.5 for Lso2). All other classes contained only 80S ribosomes with no additional factors, tRNA, or mRNA. A classification scheme is displayed in Supplemental Figure 1A.

### Structure of the native yeast Lso2-80S complex

After manual screening for ice quality, 8,600 micrographs were used for automated particle picking in Gautomatch (http://www.mrc-lmb.cam.ac.uk/kzhang/) yielding 585,801 initial particles. Upon 2D classification in RELION 3.0, 486,383 particles were selected for a consensus 3D refinement. Two rounds of 3D classification and 3D refinement and resulted in reconstructions of a low resolution 80S-Lso2 complex from 29,735 particles. This dataset was later merged with a subsequent data set, wherein, 8,400 micrographs underwent automated particle picking in Gautomatch (http://www.mrc-lmb.cam.ac.uk/kzhang/) yielding 381,233 initial particles. After 2D classification in RELION 3.0, 163,303 particles were selected for a consensus 3D refinement. 3D classification and 3D refinement and resulted in reconstructions of a low resolution 80S-Lso2 complex from 24,085 particles. The resulting merged data set of 53,820 particles underwent a consensus 3D refinement before focused sorting on the intersubunit space (A- and P site tRNA positions) resulting in one tRNA containing class of 18,869 particles and one Lso2 containing class of 34,951 particles. The Lso2 containing class underwent one final round of 3D refinement and CTF refinement resulting in an 80S-Lso2 complex reconstruction at 3.5 Å. Discarded classes contained 80S ribosomes with no additional factors, tRNA, or mRNA. A classification scheme is displayed in Supplemental Figure 1B.

### Structure of the human hibernation complex

After manual screening for ice quality, 6,145 micrographs were used for automated particle picking in Gautomatch (http://www.mrc-lmb.cam.ac.uk/kzhang/) yielding 332,890 initial particles. Upon 2D classification in RELION 3.0, 156,750 particles were selected for a consensus 3D refinement. Extensive 3D classification followed by 3D refinement and CTF refinement resulted in reconstructions of an SERBP1-eEF2-80S complex, an CCDC124-80S complex and EBP1-80S complex at 3.1 Å, 3.0 Å and 2.9 Å, respectively. A classification scheme is displayed in Supplemental Figure 4.

### Molecular models of yeast and human hibernating ribosomes

The crystal structure of the *Saccharomyces cerevisiae* (PDB: 5NDG (Prokhorova et al., 2017), the human cryo-EM structures (PDB: 6EK0 (Natchiar et al., 2017) and 4V6X (Anger et al., 2013) and the human EBP1 crystal structure (PDB: 2Q8K (Kowalinski et al., 2007a) were used as initial models to build the 80S ribosomes and EBP1, respectively. In general, the ribosome/EBP1 models were rigid body fitted into our cryo-EM maps in Chimera (Pettersen et al., 2004), followed by manually adjusting in Coot according to the densities (Emsley and Cowtan, 2004).

Due to the flexibility, the C terminal of both Lso2 and CCDC124, and the N terminal of the CCDC124 are missing in our final model, but all the other regions were *de novo* built in Coot. A homology model of the human eEF2 was generated using Swiss-Model server (Biasini et al., 2014) based on the *Sus scrofa* model (PDB: 3J7P (Voorhees et al., 2014). The human SERBP1 model was adjusted from human ribosome structure (PDB: 4V6X (Anger et al., 2013).

All the final models (Supplemental Figures 2 and 5, Supplemental Table 1) were real space refined with secondary structure restraints using the PHENIX suite (Adams et al., 2010), and the final model evaluation was performed with MolProbity (Chen et al., 2010). Maps and models were visualized and figures created with the PyMOL Molecular Graphics System (Version 1.7.4, Schrödinger, LLC) and ChimeraX (Goddard et al., 2018).

Standard model to map validations were performed according to (Brown et al., 2015a) to ensure that models are not overfitted.

## Acknowledgements

We thank S. Rieder, C. Ungewickell, A. Gilmozzi, J. Musial, and H. Sieber for excellent technical assistance. This work was supported by the German Research Council (GRK1721 and FOR1805 to R.B.) and we acknowledge support by the Center for Integrated Protein Science Munich (CiPS-M). R.Bu. is supported by a Boehringer Ingelheim Fonds PhD Fellowship and J.W. is part of the International Max Planck Research School Life Science.

## Author Contributions

J.W., R.Bu., J.C., T.B., R.B., and W.G. designed the study and wrote the manuscript. J.W. purified, biochemically characterized, prepared samples for cryo-EM, processed the data, and analyzed structures of both yeast 80S complexes with the help of J.C., T.B., R.Bu. and K.B.. R.Bu. purified, biochemically characterized, prepared samples for cryo-EM, processed the data, and analyzed the structures for human 80S complex with the help of H.K., T.B., and J.C.. J.C. built molecular models for all three structures. O.B collected cryo-EM data. J.W carried out sample preparation and splitting assays with the help of T.M.K.

