## Supplemental Material for "Structure and function of yeast Lso2 and human CCDC124 bound to hibernating ribosomes"

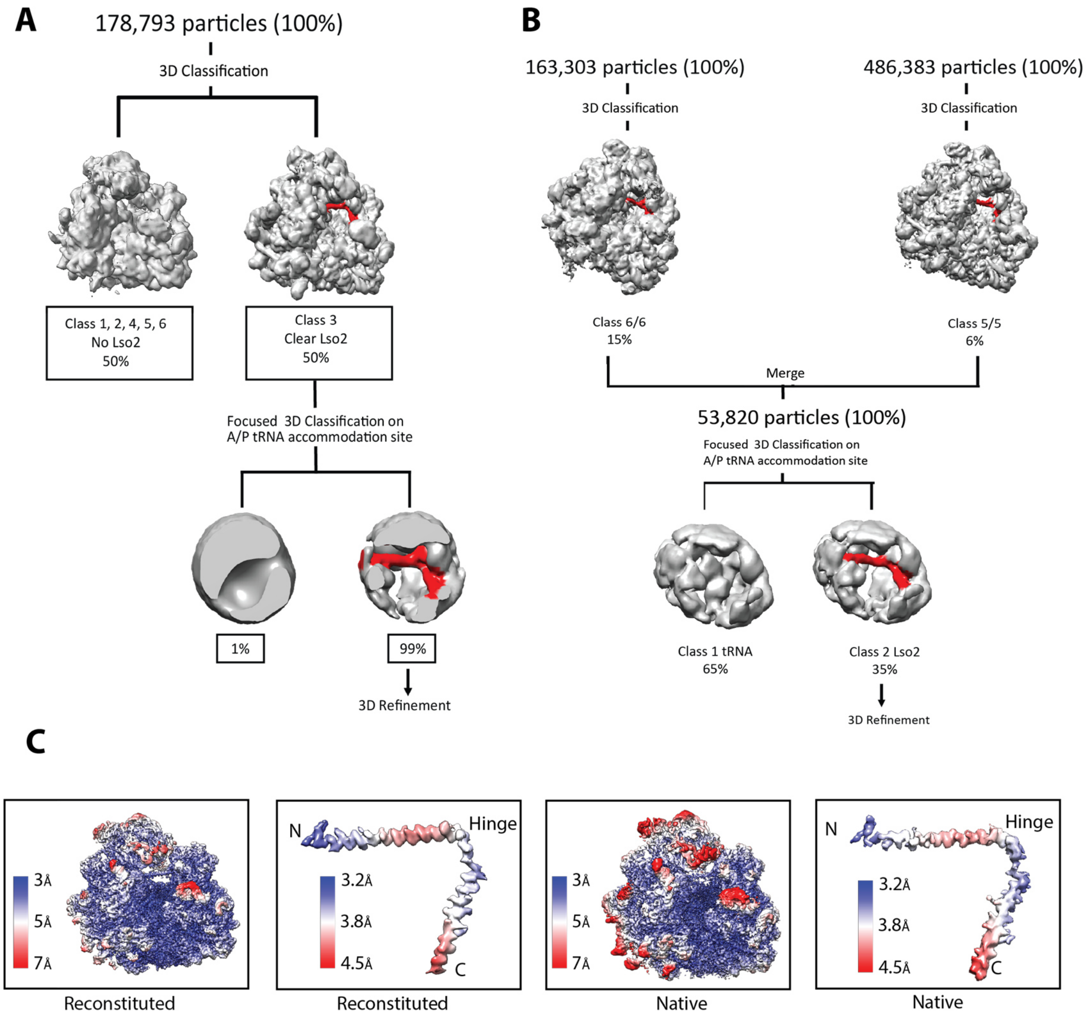


**Supplemental Figure 1:** 3D classification scheme and local resolution of Lso2-80S reconstructions.

A-B): Sorting schemes for reconstituted (A) and native (B) Lso2-80S reconstructions. For the native Lso2-80S structure particles containing Lso2 were merged from two individual collections after 3D classification due to relatively low occupancy with Lso2 (5 and 16 % of all particles, respectively). For both reconstituted and native complexes, the Lso2-containing classes were locally classified using an ellipsoid mask covering the A- and P site tRNA binding sites of the 80S. The Lso2-cotaining classes were refined to an overall resolution of 3.5 Å and 3.4 Å, respectively. C) The local resolution range for 80S ribosome and isolated Lso2 is indicated.

.

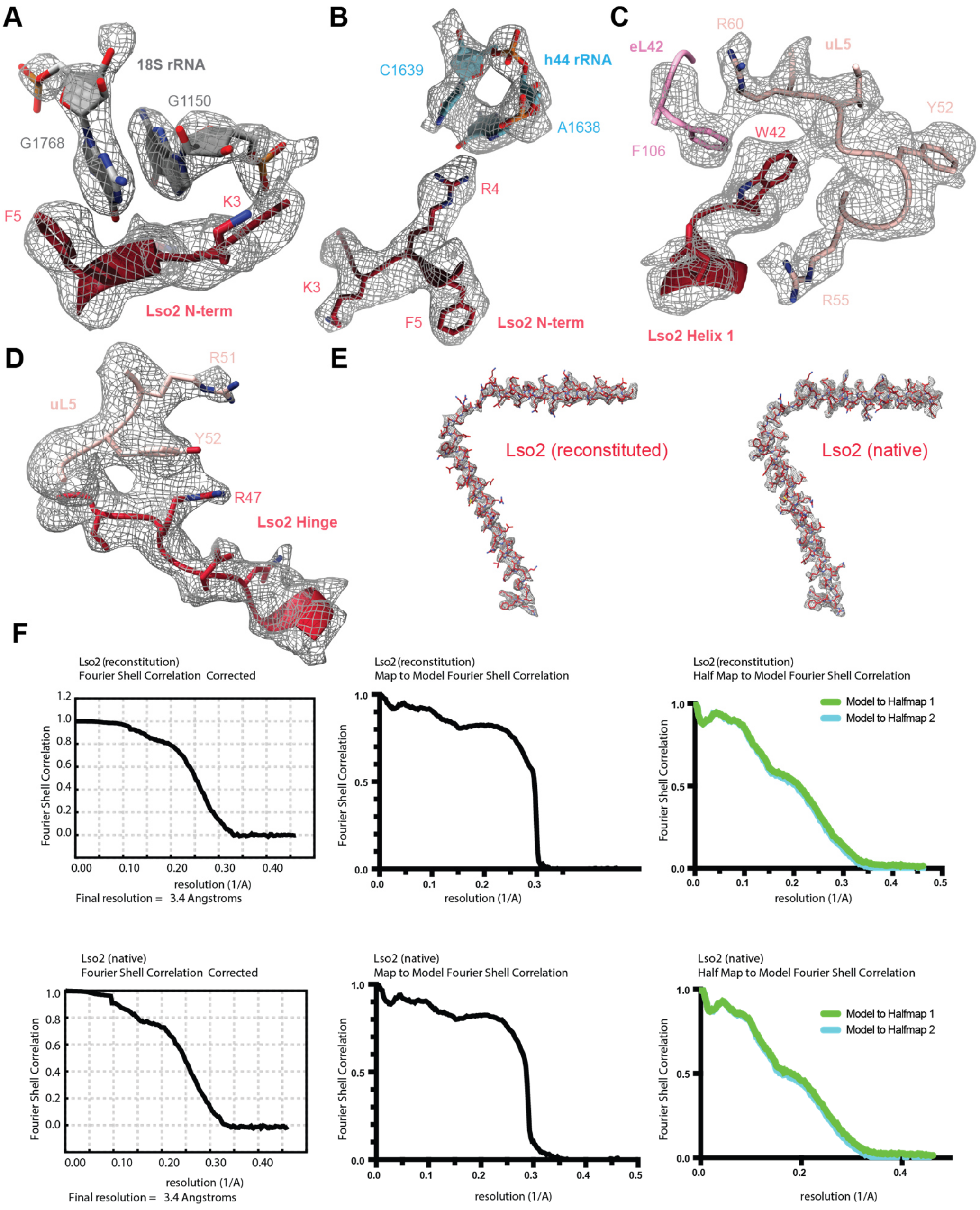


**Supplemental Figure 2:** Validation of the Lso2-80S model.
A) View highlighting the model for the N-terminus of Lso2 interacting with rRNA helix h28 (G1150) and h45 (G1768) fit into respective density (transparent grey mesh). B) View focusing on the Lso2 N-terminal R4 interacting with h44. C) View focusing on the stacking of Lso2 hinge residue W42 inside a cleft formed by LSU ribosomal proteins eL42 and uL5. D) View focusing on interactions between the hinge region of Lso2 (R47) and uL5 residues. E) Fits of the entire Lso2 model fit into isolated density. F) overall FSC curves, map to model FSC curves, and half map to model FSC curves for validation of the Lso2-80S model fitting the reconstituted and native Lso2-80S reconstructions.

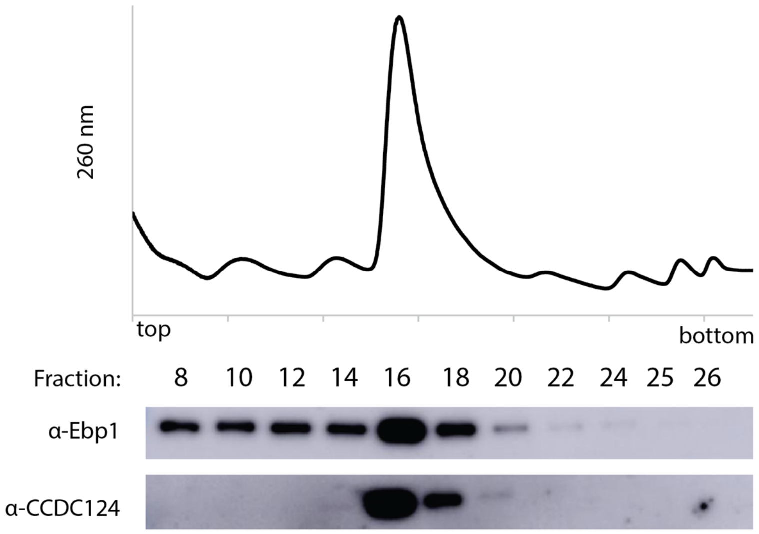


**Supplemental Figure 3**: <u>CCDC124 and EBP1 are ribosome-associated in human cell lysates.</u>
HEK293T cell lysates were fractionated through 10-40% sucrose gradients. Fractions were collected and Western blotting using antibodies against CCDC124 and EBP1 was performed. Both factors were found to associate with 80S ribosomes.

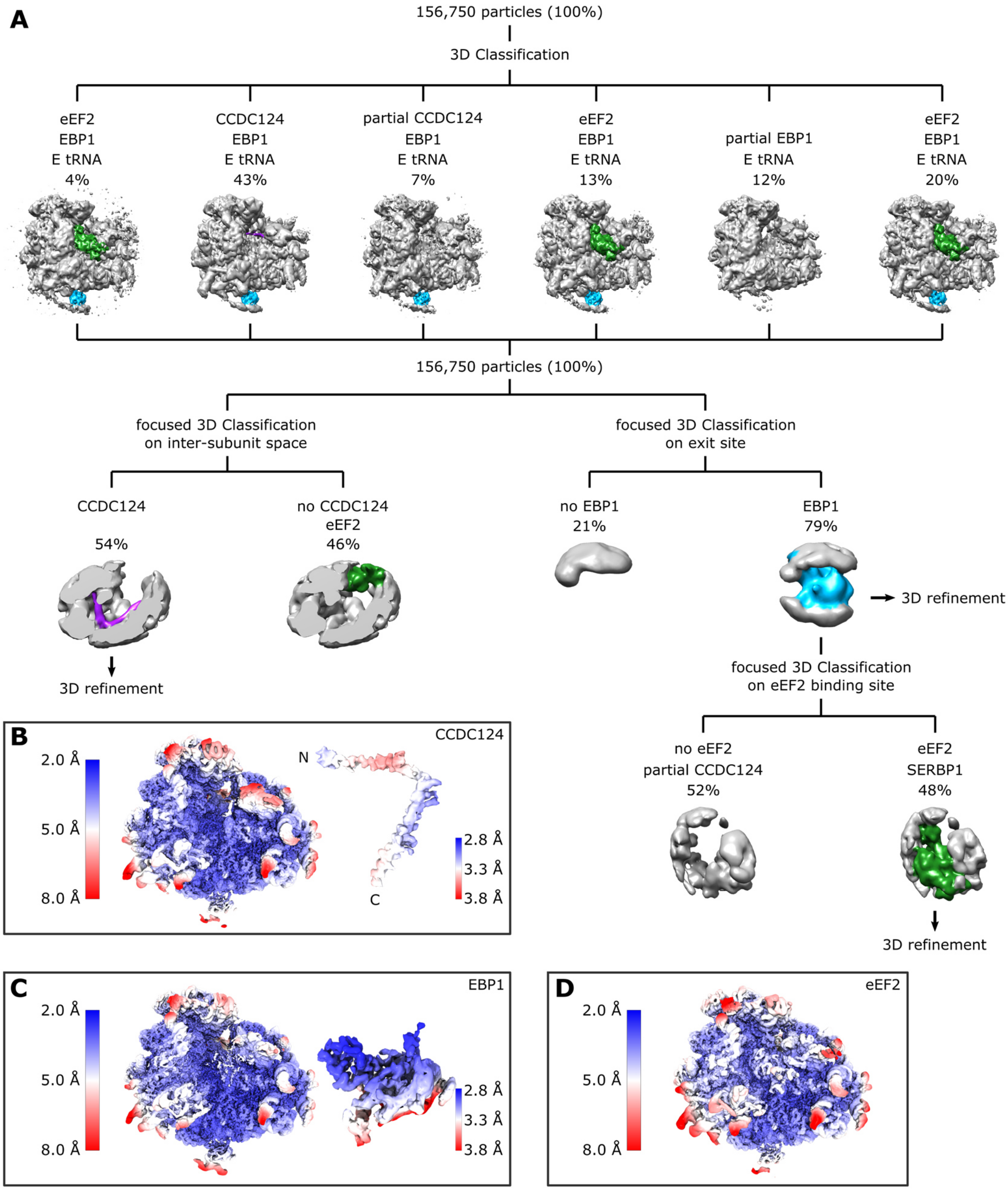


**Supplemental Figure 4:** <u>3D classification and local resolution for native human idle ribosomes.</u>

A) In a first round of 3D classification, particles were separated containing E site tRNA and EBP1 and either CCDC124 (in the non-rotated state) or SERBP1 and eEF2 (in the rotated-2) state. Independent subclassifications using a regional mask were then performed using ellipsoid masks covering the A- and P site tRNA binding sites (for CCDC124 and eEF2/SERBP1, respectively) or on the peptide exit site (for EBP1). This yielded homogenous ribosome classes containing exclusively either CCDC124 and EBP1 or SERBP1/eEF2 and EBP1. The classes displaying CCDC124-80S, SERBP1/eEF2-80S and EBP1-enriched SERBP1/eEF2-80S were refined to overall resolution of 3.0 Å, 3.1 Å and 2.9 Å. B-D) Local resolution ranges for

CCDC124-80S (B), EBP1-80S (C) and eEF2/SERPB1-80S (D), and isolated CCDC124 and EBP1 are indicated.

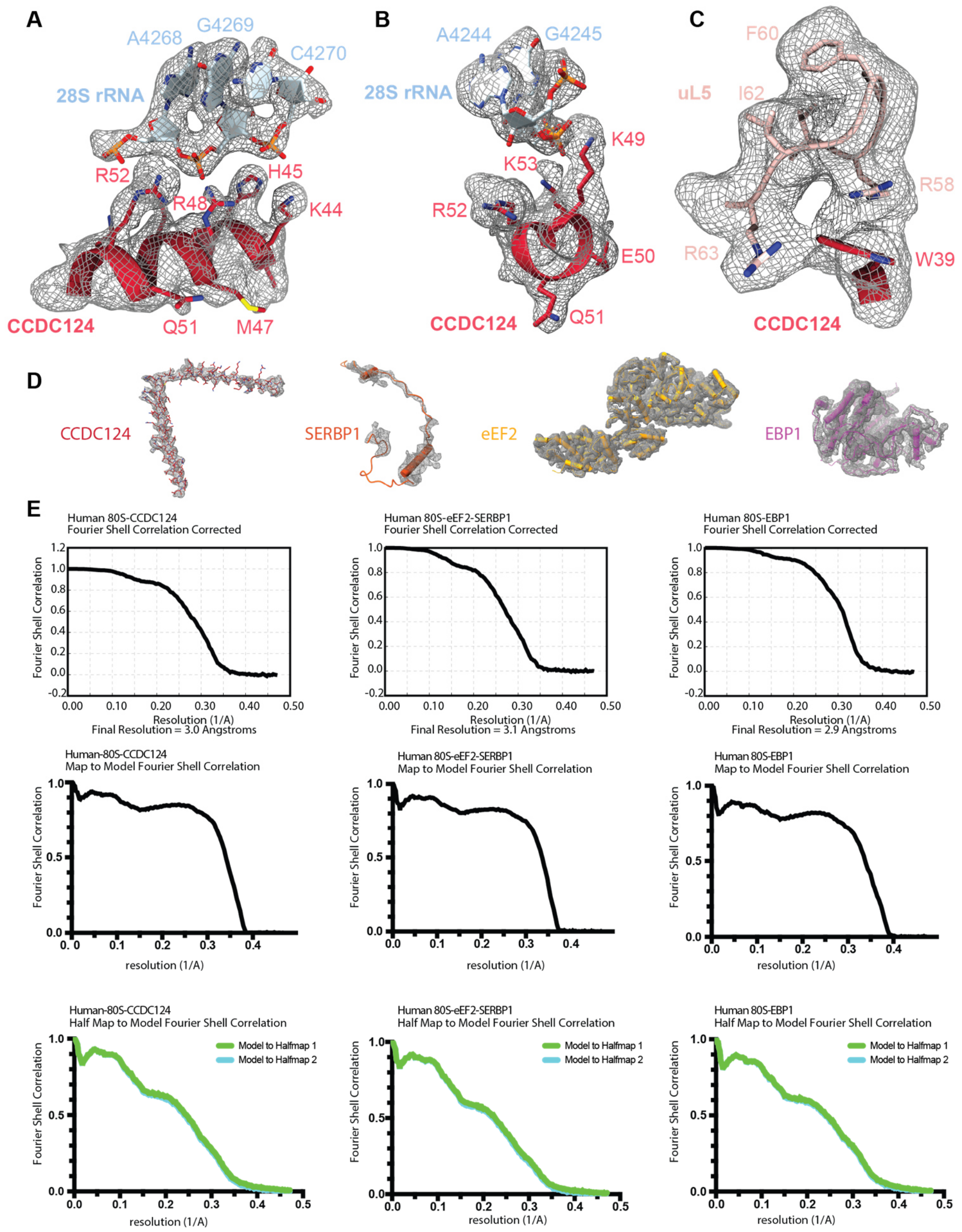


**Supplemental Figure 5:** <u>Validation of the human CCDC124-80S, SERBP1-eEF2-80S and EBP1-80S models.</u>

A-B) View highlighting the hinge region of CCDC124 interacting with H85 (A) and H84 (B) of the 28S rRNA. C) View focusing on the interaction of CCDC124 W39 stacking with residues of uL5 in a manner distinct from Lso2, wherein eL42 also participates in stabilizing Lso2 W42. D) Fits of the entire CCDC124 and EBP1 models into the respective isolated densities. E) overall FSC curves, map to model FSC curves, and half map to model FSC curves for validation of the final CCDC124-80S, eEF2/SERPB1-80S, and EBP1-80S models fitting the respective densities.

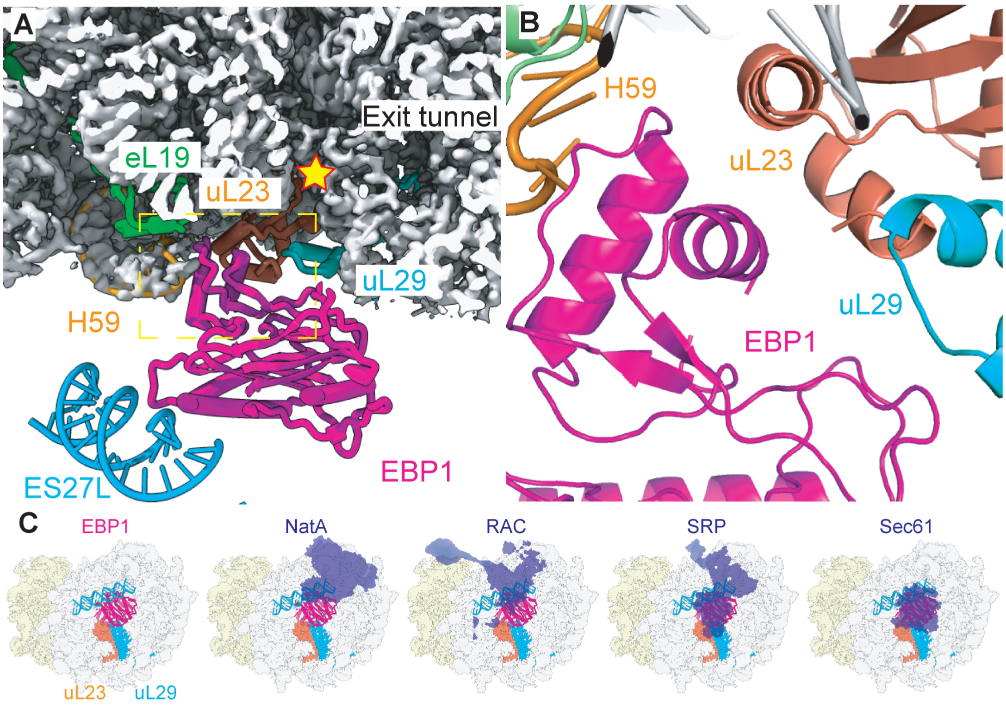


**Supplemental Figure 6:** Binding of EBP1 near tunnel exit on the 60S.

A) Close-up view showing the overall positioning of EBP1 at the peptide exit site. B) Zoom on the interaction between EBP1 and H59. C) Comparison of the EBP1 position with other exit site ligands. EBP1 binding would overlap with a majority of nascent chain interacting factors such as NatA (Knorr et al., 2019) , RAC (Zhang et al., 2014), SRP (Halic et al., 2004) or Sec61 (Becker et al., 2009; Gogala et al., 2014; Voorhees and Hegde, 2016).

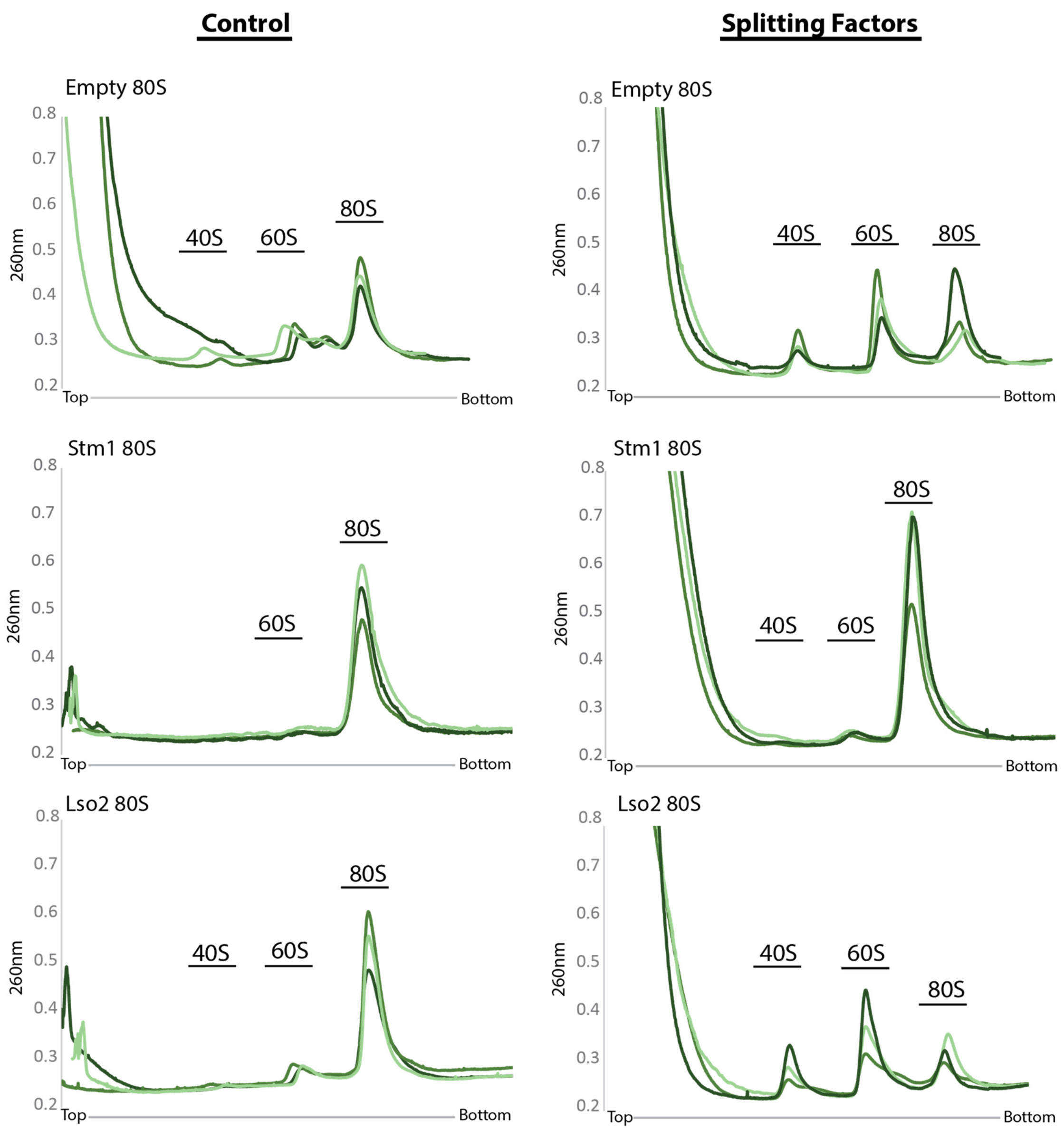


**Supplemental Figure 7:** <u>Sucrose gradient profiles for *in vitro* splitting assays</u>
UV profiles for splitting assays. Assays contain 5 pmol of ribosomes (treated with puromycin, or bound to Lso2, or bound to Stm1) and 25 pmol of factors (60S subunit anti-association factor eIF6, Dom34, Hbs1 and ABCE1). 50 µl reactions were spun through a 10-50% sucrose gradient and an absorption profile at 260 nm ($A_{260}$) was recorded. All experiments were carried out in triplicates (as indicated in different shades of green).

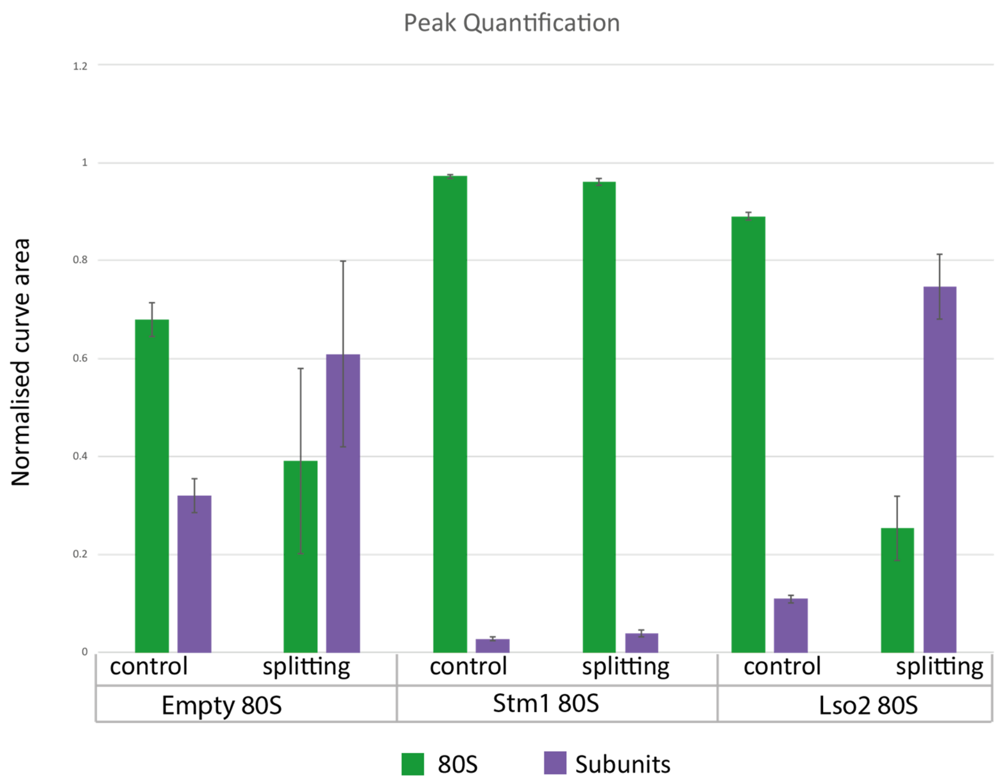


**Supplemental Figure8:** Quantification of *in vitro* splitting assays.
The relative abundance of 80S ribosomes and subunits in *in vitro* splitting assays was estimated by comparing the areas under $A_{254}$ absorption peaks from 40S, 60S and 80S subunits (calculated as described in Methods). Note, that Stm1-80S have the highest relative abundance after splitting whereas Lso2-80S show the lowest relative abundance after splitting.

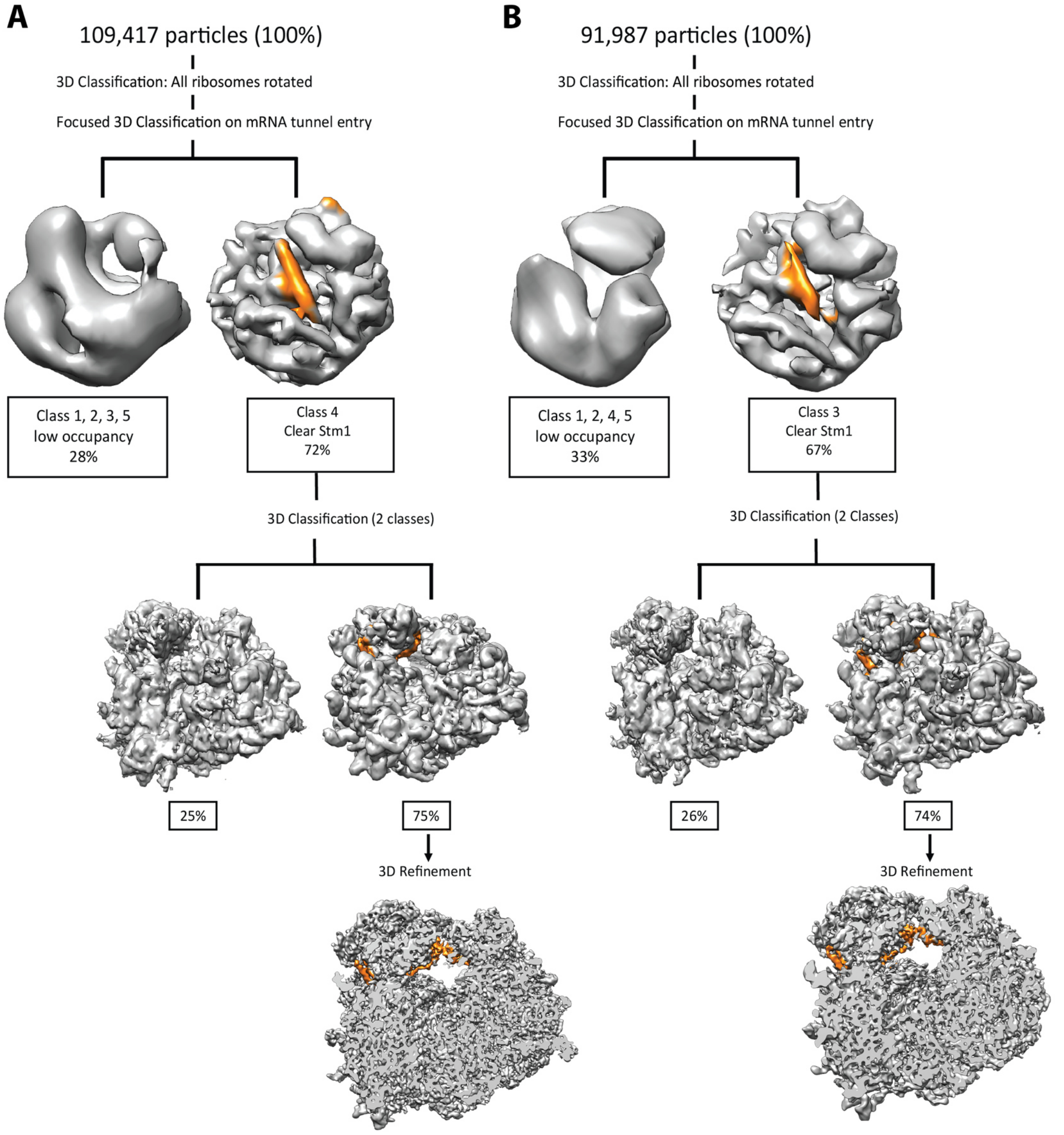


**Supplemental Figure 9:** <u>3D classification for Stm1-80S ribosomes.</u>
Stm1-80S were prepared following protocols previously reported (Ben-Shem et al., 2011) to enrich Stm1 binding and used in *in vitro* splitting reactions. Stm1-80S were collected from the sucrose gradient following *in vitro* splitting reactions with and without splitting factors (control). These 80S fractions were analyzed by cryo-EM where particles were assessed for ribosome rotational states and presence of Stm1. A) 80S ribosome population from the control splitting assay. B) 80S ribosome population remaining following incubation with splitting factors. An initial 3D classification (not shown) revealed that a vast majority of 80S are in the rotated state. These particles were further subjected to local classification using and ellipsoid mask covering the region of the mRNA channel of the 40S to classify for Stm1-containing particles (~50%) which were further refined. Stm1 density is displayed in orange.

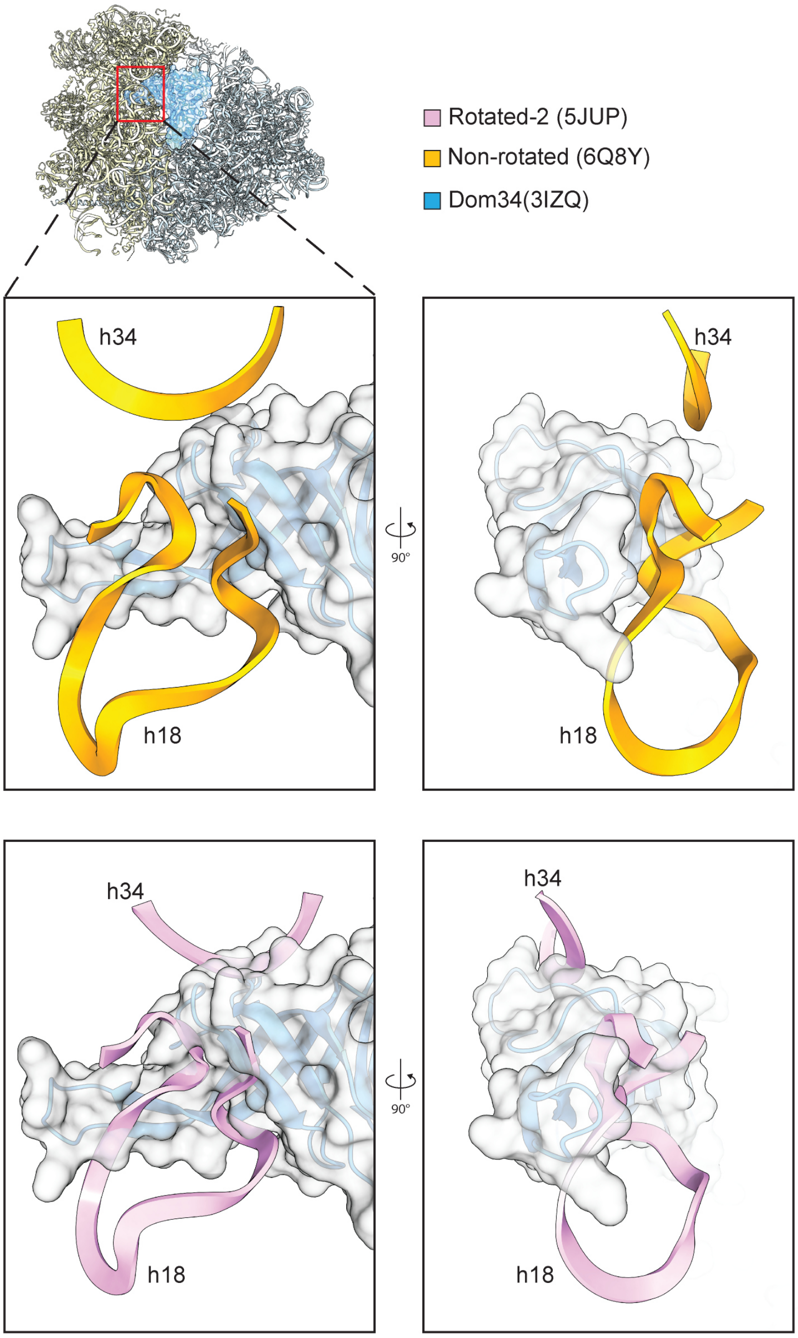


**Supplemental Figure 10:** <u>Overlay of Dom34 with 18S rRNA in rotated-2 and non-rotated states.</u>

Dom34 in complex with Hbs1 (and also ABCE1) is usually found in the non-rotated state. The structure of a yeast 80S in the non-rotated state (PDB 6Q8Y; representing Lso2-80S (Tesina et al., 2019)) was superimposed with one in the rotated-2 state (PDB 5JUP; representing Stm1-80S (Abeyrathne et al., 2016) based on the 60S subunits. We note that the Dom34 N-terminal domain (taken from PDB:3IZQ) (Becker et al., 2011) would clash with 18S rRNA helix h18 and h34 in the rotated-2 state. This indicates that the rotated (Stm1-containing) 80S ribosome is not a substrate for Dom34 and thus cannot be split by the Dom34 splitting system.

**Table 1:**

Cryo-EM data collection, refinement and validation statistics

| | Yeast ribosome +Lso2 (Native) | Yeast ribosome +Lso2 (Reconstituted) | Human ribosome +CCDC124/EBP1 | Human ribosome +eEF2/SERBP1 /EBP1 | Human ribosome +EBP1 combined |
|---|---|---|---|---|---|
| **Data collection and processing** | | | | | |
| Magnification | 75,000 | 75,000 | 75,000 | 75,000 | 75,000 |
| Voltage (kV) | 300 | 300 | 300 | 300 | 300 |
| Electron exposure (e–/Å$^2$) | 28 | 28 | 28 | 28 | 28 |
| Defocus range (μm) | -1.1 to -2.3 | -1.1 to -2.3 | -0.9 to -3.0 | -0.9 to -3.0 | -0.9 to -3.0 |
| Pixel size (Å) | 1.084 | 1.084 | 1.061 | 1.061 | 1.061 |
| Symmetry imposed | *C1* | *C1* | *C1* | *C1* | *C1* |
| Initial particle images (no.) | 649,686 | 178,793 | 332,890 | 332,890 | 332,890 |
| Final particle images (no.) | 34,951 | 88,523 | 84,429 | 72,367 | 127,706 |
| Map resolution (Å) | 3.5 | 3.4 | 3.0 | 3.1 | 2.9 |
| FSC threshold | 0.143 | 0.143 | 0.143 | 0.143 | 0.143 |
| **Refinement** | | | | | |
| Model resolution (Å) | 3.5 | 3.4 | 3.0 | 3.0 | 2.9 |
| FSC threshold | 0.5 | 0.5 | 0.5 | 0.5 | 0.5 |
| Map sharpening *B* factor (Å$^2$) | | | 90 | 80 | 90 |
| Model composition | | | | | |
| Non-hydrogen atoms | 196,487 | 196,487 | 222,325 | 228,957 | 228,587 |
| Protein residues | 11,063 | 11,063 | 12,128 | 12,949 | 12,938 |
| RNA | 5,105 | 5,105 | 5,864 | 5,876 | 5,864 |
| *B* factors (Å$^2$) | 41.80 | 62.90 | 84.50 | 110.28 | 74.33 |
| Protein | 36.11 | 56.24 | 78.95 | 109.19 | 61.94 |
| RNA | 46.37 | 68.24 | 88.93 | 111.30 | 84.74 |
| R.m.s. deviations | | | | | |
| Bond lengths (Å) | 0.0140 | 0.0106 | 0.0072 | 0.0136 | 0.0098 |
| Bond angles (°) | 1.34 | 0.93 | 0.86 | 1.07 | 0.97 |
| Validation | | | | | |
| MolProbity score | 2.14 | 2.28 | 2.01 | 2.16 | 1.92 |
| Clashscore | 6.58 | 14.86 | 10.07 | 13.73 | 7.94 |
| Poor rotamers (%) | 1.92 | 0.64 | 0.46 | 0.68 | 0.67 |
| Ramachandran plot | | | | | |
| Favored (%) | 89.59 | 87.88 | 92.00 | 91.32 | 91.86 |
| Allowed (%) | 10.01 | 11.64 | 7.83 | 8.49 | 7.93 |
| Disallowed (%) | 0.40 | 0.48 | 0.17 | 0.19 | 0.21 |
